# Genomic signatures of host-specific selection in a parasitic plant

**DOI:** 10.1101/2022.02.01.478712

**Authors:** Emily S. Bellis, Clara S. von Münchow, Alan Kronberger, Calvins O. Odero, Elizabeth A. Kelly, Tian Xia, Xiuzhen Huang, Susann Wicke, Steven M. Runo, Claude W. dePamphilis, Jesse R. Lasky

## Abstract

**Premise:** Parasitic plants and their hosts are model systems for studying genetic variation in species interactions across environments. The parasitic plant *Striga hermonthica* (witchweed) attacks a range of cereal crop hosts in Africa and exhibits substantial variation in performance on different host species. Some of this variation is due to local adaptation, but the genetic basis of specialization on certain hosts is unknown.

**Methods:** To identify genomic regions that are strongly differentiated between parasites attacking different host species, we present an alignment-free analysis of *S. hermonthica* population diversity using whole genome sequencing (WGS) data for 68 individuals from western Kenya. We validate our findings with germination experiments and analyses based on a *de novo* assembled draft genome.

**Results:** Reference-free and reference-based analyses suggest that only a small portion of the *S. hermonthica* genome is strongly differentiated by host species in populations from western Kenya. Analysis of host-associated *k*-mers implicated genes involved in development of the parasite haustorium (a specialized structure used to establish vascular connections with host roots) and a potential role of chemocyanins in molecular host-parasitic plant interactions. Conversely, no phenotypic or genomic evidence was observed suggesting host-specific selection on parasite response to strigolactones, hormones exuded by host roots and required for parasite germination.

**Conclusions:** This study demonstrates the utility of WGS for plant species with large, complex genomes and no available reference. Contrasting with theory emphasizing the role of early recognition loci for host specificity, our findings support host-specific selection on later interaction stages, recurring each generation after homogenizing gene flow.

## INTRODUCTION

Characterizing the genomic basis of adaptation to local biotic environments is a key challenge for evolutionary ecology (Ebert and Fields, 2020). Parasites and mutualists exert strong influences on host fitness, and may even constitute the predominant selective pressure shaping patterns of local adaptation in some systems (Fumagalli et al., 2011; Castellano et al., 2019). Many host-parasite and host-mutualist systems involve a complex multi-step infection process including many stages of interaction between host and symbiont derived molecules (Hall et al., 2017). An outstanding question is whether adaptation to local biotic environments occurs most often via selection on genes involved during initial infection stages, or whether genetic variation at later stages of the interaction is also frequently maintained.

The expectation from theoretical studies is that initial recognition loci are more likely than downstream effector loci to contribute to host genotype by parasite genotype interactions (G_H_ x G_P_) and correspondingly, local adaptation (Nuismer and Dybdahl, 2016). The first prediction, that recognition loci contribute more to G_H_ x G_P_, is supported by empirical studies of the waterflea *Daphnia magna* and its bacterial parasite *Pasteuria ramosa* (Hall et al., 2017). In this system, most of the genetic variance in parasite infection was associated with a single major effect QTL linked to the early stage of parasite attachment (Hall et al., 2019). In contrast, many different QTLs of smaller effect were associated with later stages, highlighting the potential for independent evolution of traits involved in different stages of the infection process (Hall et al., 2019). Supporting the second prediction that selection on recognition traits often underlies local adaptation (Nuismer and Dybdahl, 2016), studies of plant pathosystems have revealed reciprocal coevolutionary selection on host resistance (R) genes and parasite avirulence genes, for example in the flax-flax rust system (Ravensdale et al., 2011; Thrall et al., 2012). However, a high degree of genotype specificity has also been observed for many host-parasite systems characterized by more quantitative mechanisms of resistance (Poland et al., 2009). For host-parasite interactions characterized by quantitative genetic architectures, we still know little regarding the genetic basis of local adaptation in natural populations and the extent to which initial vs. later infection stages contribute to G_H_ x G_P_.

An emerging model system for studying spatial pattern and process in coevolutionary genomics is the parasitic plant *Striga hermonthica* (family Orobanchaceae) and its cereal hosts. In contrast to *Striga gesnerioides,* which parasitizes cowpea via a qualitative gene-for-gene mechanism, natural variation in host resistance to *S. hermonthica* is highly polygenic (Li and Timko, 2009; Timko et al., 2012) with at least one large effect locus (Gobena et al., 2017). *Striga hermonthica* parasitizes grass hosts including sorghum, maize, rice, and millets and is one of the greatest biotic constraints to food security in Africa (Ejeta, 2007; Spallek et al., 2013; Savary et al., 2019). An individual *S. hermonthica* plant can produce thousands of seeds that are speculated survive in the soil for a decade or more under optimal conditions (Bebawi *et al*. 1984; but see Gbèhounou *et al*. 2003).

A great deal has been learned in the past decades regarding the mechanisms through which *Striga* parasitizes diverse hosts. To germinate, parasite seeds must detect strigolactones (SLs), hormones exuded from host roots under nutrient-deficient conditions that also stimulate host interactions with beneficial mycorrhizal fungi (Akiyama et al., 2005). Perception of SLs is mediated through binding to paralogs of KARRIKIN INSENSITIVE 2 (KAI2), known as HYPOSENSITIVE TO LIGHT (HTL) receptors. SL receptors of the *KAI2d* clade rapidly expanded and diversified during the transition to parasitism in the Orobanchaceae, a plant family that includes thousands of mostly root parasitic species (Conn et al., 2015). For example, the ∼600 Mb genome of *Striga asiatica* contains 21 *KAI2* genes, many of which occur as tandem duplications (Yoshida et al., 2019). *Striga hermonthica* may possess 13 or more *KAI2* paralogs (Nelson, 2021), including 11 for which the binding affinity for diverse SLs has been extensively characterized (Toh et al., 2015; Tsuchiya et al., 2015).

Following SL perception and parasite germination, host-derived phenolic compounds induce formation of the haustorium, the specialized multicellular feeding structure used by parasitic plants to infest host tissues (Cui et al., 2018). Intrusive cells of the haustoria invade host tissues to form direct connections with host vasculature (Masumoto et al., 2021). Water, nutrients, and other molecules including mRNA (Kim et al., 2014), small RNA (Shahid et al., 2018), DNA (Yang et al., 2019), and proteins (Liu et al., 2020; Shen et al., 2020) can be directly transferred across haustorial connections. In addition to low germination stimulation (Dayou et al., 2021; Mallu et al., 2021), post-germination host resistance to *Striga hermonthica* can occur at later interaction stages through induction of an intense hypersensitive response, formation of a mechanical barrier, or failure of the parasite to form vascular connections (Mbuvi et al., 2017; Mutinda et al., 2018; Kavuluko et al., 2020). Much of the genetic variation in these mechanisms of resistance may result from host populations’ local adaptation to parasitism across the range of *Striga hermonthica* (Bellis et al., 2020).

Conversely, many parasite populations may also be adapted to their local host communities. Across the geographic range of *S. hermonthica* in Africa, there exists considerable variation in host crop communities, and regional abundance of a particular host species is associated with high *Striga* performance on that crop (Bellis et al., 2021). Given the low germination of millet- and sorghum-specific *S. hermonthica* populations in response to root exudates from the alternate host (Parker and Reid, 1979), it is possible that at least some of this host specialization results from natural selection on SL perception. Host-specific germination by *Striga* genotypes could potentially result from duplication of different *KAI2* paralogs and neofunctionalization to fine-tune the response to different SLs, with host community heterogeneity maintaining presence/absence variation (Nelson, 2021). In contrast, if parasite local adaptation involves later interaction stages, ‘core parasitism genes’ implicated in haustorium development in previous studies (Yang et al., 2014) could be the targets of host-specific selection.

Extensive genetic and germplasm resources for host species have enabled broad-scale studies of sorghum local adaptation to *S. hermonthica* parasitism (Bellis et al., 2020), but understanding the genetic basis of reciprocal adaptation is challenged by a paucity of genomic data for parasites. At the population-scale, restriction-site associated DNA sequencing (RAD- Seq) and analysis of polymorphism in transcriptomes have begun to shed light on population level diversity in *S. hermonthica* parasites (Unachukwu *et al*. 2017; Lopez *et al*. 2019). However, reduced representation approaches pose difficulties if only a fraction of host-associated genome diversity is tagged by RAD-Seq markers (Lowry et al., 2017) or (in the case of transcriptomes) if genes under selection are not expressed in sequenced tissues. Like other parasitic angiosperms, *S. hermonthica*, a diploid species with *n* = 19 or 20 (Iwo et al., 1993; Aigbokhan et al., 1998), is characterized by a larger genome than non-parasitic relatives (Lyko and Wicke, 2021), with estimated genome size of ∼1 Gb (1C = 0.9 Gb; Yoshida *et al*. 2010) or greater (1C = 1.4 Gb; Estep *et al*. 2012). Several genome assembly projects are currently underway, but high heterozygosity, obligate outcrossing, and a large genome pose substantial challenges for assembly, and a publicly available reference for *S. hermonthica* is not currently available (but see Qiu et al., 2022 for another genome assembly that may be available soon). Here, we provide a reference-free analysis of population-scale diversity in *S. hermonthica,* based on whole genome sequencing (WGS) data. Using a unique alignment-free bioinformatic approach, we identify genetic variation associated with host-specific parasitism in natural populations and investigate signatures of selection surrounding candidate loci. Based on these findings, we evaluate the hypothesis that adaptation to local host populations results from selection on genes involved in early stages of the interaction (SL perception) against the alternative that selection on genes involved in later stages is primarily responsible.

## MATERIALS AND METHODS

### Sample Collection

Seeds and leaf tissue were collected from *S. hermonthica* individuals in July 2018 from six locations in western Kenya (Fig. 1A), which are location-matched with previously collected specimens in herbaria. Two plots with *S. hermonthica* parasitizing different host species were sampled at each location, chosen as close as possible and in most cases, from nearby plots on the same farm (i.e. within 15 meters). In one location (Mumias), we were not able to find a parasite population on another host, so two maize- parasitizing populations were sampled. Per plot, twelve *S. hermonthica* individuals were chosen haphazardly, with effort taken to sample individuals distributed evenly throughout the plot.

**Figure 1.**
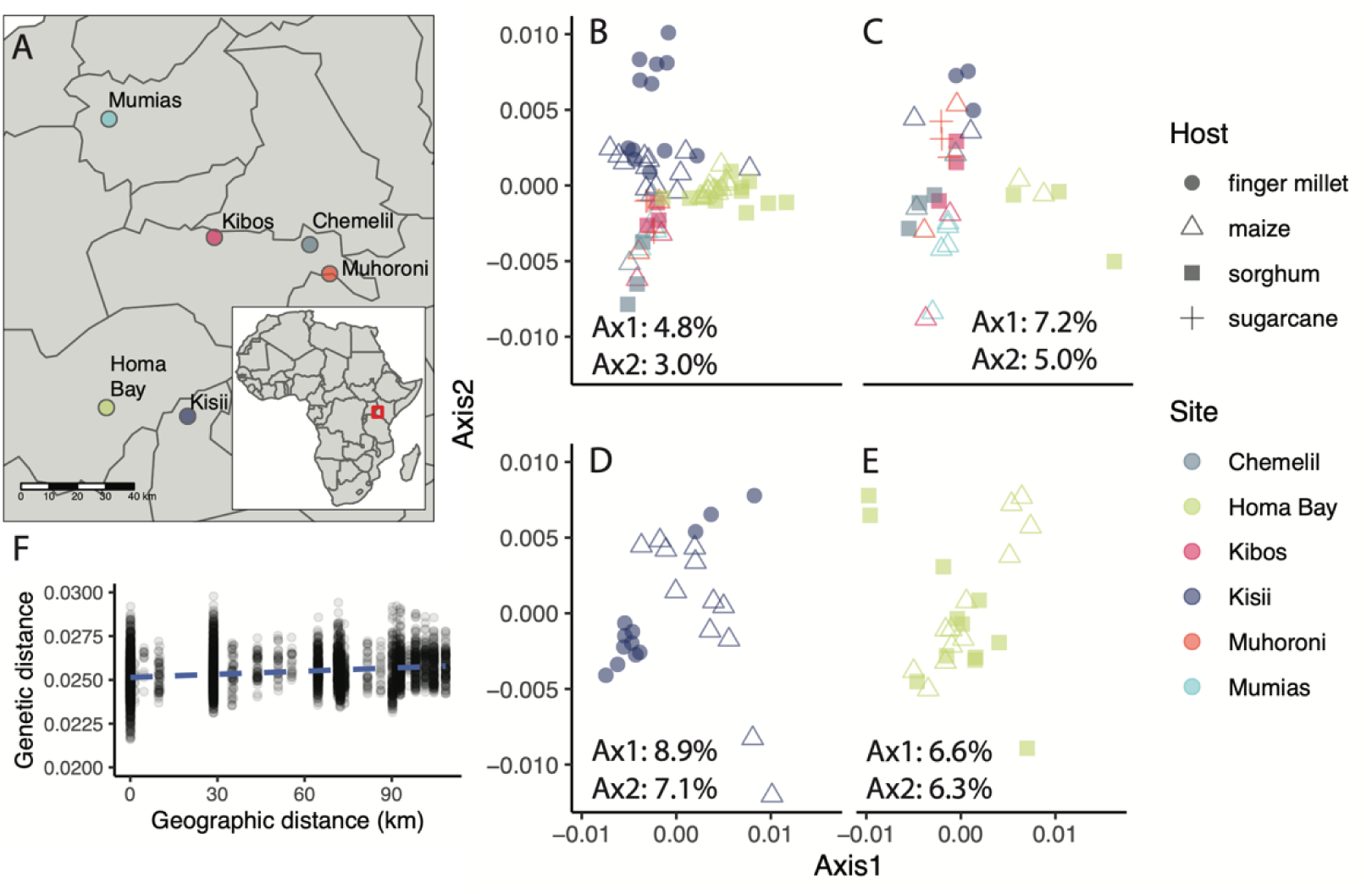
Population genomics of *S. hermonthica* from western Kenya. A) Map of the six sampling locations. (B-E) Principal Coordinates Analysis (PCoA) based on *k-*mer-derived genomic distances, performed separately for B) all sampled individuals (*n* = 68), C) five individuals per location (*n* = 30), D) individuals from Kisii (*n* = 24), and E) individuals from Homa Bay (*n* = 24). Relative proportion of variability explained by the first and second principal coordinate axes is indicated for each analysis. F) Genetic vs. geographic distance. Genetic distance was based on 31-mers. The dashed blue line indicates expectations from the best fit line (y = 0.000013 * km + 0.0246; *R^2^* = 0.05) for *n* = 4,556 pairwise comparisons among different individuals.

Individuals were photographed before collection, and images for representative individuals at each site were uploaded to iNaturalist. Three to four leaves per individual were sampled directly into silica gel, before collection of the whole individual into separate paper bags. Plants were dried in paper bags before harvesting and manual cleaning of seeds. Cleaned seeds were shipped to Penn State for germination rate experiments and stored in individual 2 mL microcentrifuge tubes at room temperature prior to experiments. Five voucher specimens (ESB collection numbers 2018.1 to 2018.5) were deposited in the collection of the East African Herbarium (EA).

### Whole genome sequencing

For samples collected in 2018, whole genome sequencing was performed for a subset of 68 *S. hermonthica* individuals. This included all 24 individuals collected from adjacent plots of finger millet and maize in Kisii and all 24 individuals collected from adjacent plots of sorghum and maize in Homa Bay. We hypothesized that for each of these locations, parasites sampled from different hosts would show overall high similarity across much of the genome due to high rates of gene flow among neighboring plots. However, we expected to identify loci under strong host-specific selection as differentiated against this backdrop. To interpret patterns of host-specific differentiation for the Homa Bay and Kisii populations in the context of broader population genetic diversity in western Kenya, we also sequenced DNA from five individuals (two or three per host species) from four additional locations. DNA was extracted from silica-dried leaf tissue in the USDA APHIS quarantine facility at the Pennsylvania State University using the E.Z.N.A. Plant DNA DS Mini Kit (Omega Bio-tek, Norcross, Georgia, USA) according to the manufacturer’s protocol. Genomic library preparation and paired-end 150 bp sequencing was carried out by the Texas A&M AgriLife Genomics and Bioinformatics Service, on a single lane of a NovaSeq 6000 S4 flow cell.

### Population structure

To determine whether parasites from the same location showed the expected pattern of low genetic differentiation between different hosts, we followed a reference-free approach to evaluate population genomic patterns among sequenced samples. Raw reads that could be classified as plant-derived were identified using Kraken 2 (Wood et al., 2019), based on a custom database built from the complete set of plant genomes and proteins in the NCBI RefSeq collection, sequences from 472 Mbp of the *Striga asiatica* genome (Yoshida et al., 2019) and *S. hermonthica* transcriptome sequences (build StHeBC4) from the Parasitic Plant Genome Project II (Yang et al., 2014). Classified sequences were trimmed using BBduk from BBTools (Bushnell, n.d.), removing sequence on the ends of reads with low quality (qtrim=rl trimq=20 minlen=50) or 3’ matches to adapters (k=23 mink=11 hdist=1 tpe tbo ktrim=r). To reduce bias associated with differences in per-sample read depth, reads were downsampled to 5.2 Gbp with BBTools Reformat.

*Mash*, a dimensionality reduction technique based on the MinHash algorithm, was used to estimate genetic distance between samples based on resulting read sets (Ondov et al., 2016). *Mash* previously showed improved performance compared to alignment-based methods for estimating pairwise genetic distance for polyploid plant genomes using simulated and real data (VanWallendael and Alvarez, 2020). We used a *k-*mer size of 31, removing *k*-mers with less than 2 copies but increasing the sketch size to 1 x 10^7^ to account for a larger volume of input data. Principal Coordinates Analysis was performed in R version 4.0 with the pcoa function of the ‘ape’ package (Paradis and Schliep, 2019). A shorter *k-*mer size (*k*=21) was also tested but did not alter clustering patterns in PCoA. Correlation between the genetic distance matrix and the geographic distance matrices, calculated with ‘geodist’, was determined using a Mantel test (Padgham and Sumner, 2021).

If contamination is present in our Kraken2 plant genome database, our reference-free analysis of population diversity could be biased by contaminant sequences. Therefore, we also included an analysis of population structure based on alignment to our assembled reference genome (see “Validation of *k*-mer based approaches”). After removing scaffolds shorter than 500 bp from the reference, high quality sequences from each *S. hermonthica* individual were mapped using BWA-MEM v0.7.17 (Li, 2013). Sequences mapping with quality less than 20 were excluded using SAMtools v1.10 (Li et al., 2009). Pairwise genetic relationships were calculated from genotype probabilities using *ngsDist* (Vieira et al., 2016). Genotype posterior probabilities were assigned using ANGSD v0.935 (Korneliussen et al., 2014), ignoring sites with a minimum coverage less than 50 or greater than 500 across all 68 individuals and contigs less than 2.5 kb.

### Host-specific differentiation

We next sought to identify particular genomic regions differentiated between parasites growing on different hosts. This analysis targeted parasite populations from Homa Bay or Kisii, for which we sequenced DNA from parasites for two different hosts on the same farm from immediately adjacent plots. For the Kisii population, the dataset included individuals from finger millet and maize, whereas for the Homa Bay population the dataset included parasites from sorghum and maize (*n* = 12 from each host species; 48 individuals total). Counts for *k*-mers of length 31 were summarized across sequenced individuals using HAWK; 31-mers were used as the longest *k*-mer that can be represented efficiently on a 64-bit machine (Rahman et al., 2018). At each *k*-mer we considered two allelic states (present or absent) where the *k*-mer was marked as present in an individual if it was counted at least twice or absent if it was not observed at all. A single biallelic SNP in otherwise invariant 31 bp of sequence, for example, would be represented by two distinct 31-mers. Note that *k-*mers which only appear once in samples are filtered by HAWK by default (Rahman et al., 2018). Allelic states were used to calculate the fixation index, *G_ST_*(Nei and Chesser, 1983), for each *k*-mer using custom Python scripts (https://github.com/em-bellis/StrigaWGS). *G_ST_*is a generalization of the widely used fixation index *F_ST_* applicable to non-diploid loci (Nei, 1973); for the purposes of our *k*-mer analysis, we consider loci as haploid rather than diploid, since only presence or absence of the *k*-mer is scored. While we did not remove *k*-mers with excessively high counts (likely derived from repetitive genomic sequences) these are not likely to be identified as *G_ST_*outliers since the *k*-mer would need to have a count of zero in most samples from one population but not the other.

To gather functional information for host-associated *k*-mers, 31*-*mers with *G_ST_* above 0.5 were extracted and assembled into longer contigs using ABYSS 2.0.2 (Jackman et al., 2017), specifying a *k-*mer length of 29 for assembly. The *k*-mer length for assembly was chosen to facilitate assembly of relatively longer contigs (minimum expected length of 59 bp, if formed by two 29-mers overlapping with a SNP site on either side). To evaluate significance of the chosen *G_ST_* threshold, we carried out permutation analyses for Homa Bay and Kisii populations separately. For each table (unique *k*-mers in rows and individuals in columns), the host label for each individual was randomly permutated, and *G_ST_* values for each *k-*mer recalculated. For each permutation, we then calculated the proportion of *k*-mers with *G_ST_* > 0.5, to estimate the proportion of ‘outlier’ *k*-mers at our chosen *G_ST_* threshold expected by chance. Assembled contigs were then queried against contigs from two published *S. hermonthica* transcriptome assemblies using BLAST optimized for short sequences (blastn-short). Transcriptome assemblies in our BLAST database included StHeBC4 (265,694 scaffolds covering 369.7 Mb) from the Parasitic Plant Genome Project (Westwood et al., 2012) and Sh14v2 (81,559 scaffolds covering 83.9 Mb) from Yoshida *et al*. (2019). Annotations are based on the top hit from the Sh14v2 transcriptome.

To further investigate genomic regions associated with host-specific differentiation, we mapped cleaned sample reads to mRNA reference sequences, following the strategy from Therkildsen and Palumbi (2017). Mapping to a transcriptome reference has the potential to introduce errors in SNP calling due to intron/exon boundaries and the short length of transcripts, so we focused this analysis on a small set of loci for which alignments could be manually inspected including three transcripts with potential functions in haustorium development (StHeBC4_h_c11261_g0_i1, StHeBC4_p_c12587_g2_i1, StHeBC4_u_c12903_g27039_i4; see Results). The reference also included transcript sequences for a set of 11 previously characterized *S. hermonthica* strigolactone receptors [GenBank accession numbers KR013121 - KR13131] (Tsuchiya et al., 2015). High quality, contaminant-filtered reads were mapped to the reference transcriptome using BWA-MEM (Li, 2013), and alignments with low quality were removed using SAMtools view (-q 20) (Li et al., 2009). Allele frequencies for each site in the reference transcriptome were estimated based on genotype likelihoods using ANGSD, ignoring low quality bases (-minQ 25), allowing reads for which only one end mapped (- only_proper_pairs 0), and assuming known major and minor alleles but accounting for uncertainty of the minor allele (-doMaf 3) (Kim et al., 2011; Korneliussen et al., 2014). Sites with information for fewer than nine of twelve individuals in the population were excluded, and the difference in estimated allele frequency between parasite populations on different hosts was visualized with R version 4.0 (R Core Team, 2020). This strategy was used to filter false SNP calls due to errors in mapping DNA-derived reads to a transcriptome reference, since these SNPs should be observed as a fixed difference from the reference that occurs in both populations. We further investigated genes with potential structural variation by extracting reads aligned to the transcript reference and their unmapped pairs and reassembling them with ABYSS (k=51) (Jackman et al., 2017).

### Validation of k-mer-based approaches

To investigate patterns of selection surrounding putative host-associated loci and validate findings from reference-free analyses, we also mapped reads generated for *S. hermonthica* to a draft reference assembled specimen, grown *ex situ* from seeds collected on maize in the Irimbi district of Southern Uganda (specimens voucher deposited at MSUN). DNA was extracted from developing leaves and inflorescences of one individual using a modified 1x CTAB-protocol with subsequent PEG-8000 precipitation and purification (Wicke et al., 2016). Genomic libraries were sequenced on an Illumina HiSeq 2000 in 101 bp paired-end mode at Eurofins GATC Biotech GmbH (Constance, Germany). Additional data were generated to a targeted depth of 110X using the HiSeq 2500 platform using the HiSeq SBS Kit v4 at Eurofins GATC Biotech GmbH, for which DNA from the original extract was subjected to Φ29-polymerase based whole-genome amplification. Whole genome-amplified DNA was size-selected for >20 kb fragments on 1% low-melting point agarose and purified using an agarose digest-based purification with subsequent ethanol/sodium acetate precipitation (Wicke et al., 2013). For the final assembly, we employed Trimmomatic v0.36 (Bolger et al., 2014) to remove adapters and retain only sequences longer than 36 bp with an average per-base quality above 15 (“ILLUMINACLIP:TruSeq3-PE.fa": 2:30:10 SLIDINGWINDOW:4:15 MINLEN:36). The quality-trimmed data were assembled using SPAdes v3.10.1 with *k-*mer sizes of 21, 33, 55, and 77 (Bankevich et al., 2012). Assembly quality was ascertained using Quast v4.5 (Gurevich et al., 2013) with default parameters for eukaryotes. The resulting assembled contigs were contaminant-filtered, for which we ran a nucleotide BLAST search of all contigs against the non-redundant nucleotide database (access date: 10.07.2017) using BLAST+ v2.6 with an e-value of 1E10^-4^. Only contigs with the three best hits matching to green land plants (Viridiplantae) were retained.

After removing scaffolds shorter than 500 bp from the reference, high quality sequences from each *S. hermonthica* individual were mapped using BWA-MEM v0.7.17 (Li, 2013). Sequences mapping with quality less than 20 were excluded using SAMtools v1.10 (Li et al., 2009). Tajima’s D was calculated in non-overlapping windows of 1-kb using ANGSD v0.935 based on the folded site frequency spectrum and including reads where only one end mapped (- only_proper_pairs 0), to account for the highly fragmented nature of the assembly (Korneliussen et al., 2014). *F_ST_* was also calculated in non-overlapping windows with ANGSD, using the SAMtools method for calculating genotype likelihoods. Genome-wide mean values of Tajima’s D were determined by fitting an intercept-only linear mixed model to window estimates of *F_ST_* or Tajima’s D, including a random effect of ‘contig’ to account for increased correlation among measurements from nearby genomic regions, with the R package lme4 (Bates et al., 2015). An empirical *P-*value for Tajima’s D for the contig containing the focal chemocyanin was calculated based on the number of 10,000 randomly sampled windows of size matching the assembled length of the contig (1-kb) with values more extreme than the observed value.

To validate the presence/absence polymorphism for the focal chemocyanin gene, we performed PCR using primers designed from the reassembled ‘finger millet’ allele (Primer Set A: 5’-AAGATTGCGGTTACCACCAG-3’ and 5’-TCTCGATCCTTTTGGAATGG-3’) and the transcript reference (Primer Set B: 5’- CAGGAGCAAGTAGAGTAGAGCA-3’ and 5’- TGGGGAAAGAGGTAGTGCAA-3’). PCR was performed with DreamTaq DNA Polymerase 2x Mastermix (ThermoFisher, Waltham, Massachusetts, USA) under the following cycling conditions: 95°C for 3 min; 30 cycles of 95°C (30 s), 50.3°C (30 s), 72°C (60 s); 72°C for 5 min.

### Germination experiments

Seed germination was assayed in the USDA-APHIS- permitted quarantine lab at Pennsylvania State University (permit no. P526P-21-04540). A detailed step-by-step protocol is available from protocols.io repository (Bellis and Kelly, 2019), following the modifications for testing seeds collected from individual plants. Briefly, seeds were surface sterilized for 10 minutes in 1.5 mL microcentrifuge tubes with a 0.5% sodium hypochlorite solution before preconditioning for 12 days at 30°C in separate wells of foil-wrapped 12-well culture plates, with three replicates per unique germination stimulant and parasite genotype combination. Germination counts were performed 3 days after addition of germination stimulants. Tested germination stimulants included (+)5-deoxystrigol (Olchemim, Olomouc, Czech Republic; CAS: 151716-18-6) or (±)orobanchol (Olchemim; CAS: 220493-64-1) at 0.01 µM and (±)-GR24 (Chempep, Wellington, Florida, USA; CAS: 76974-79-3) at 0.2 µM. GR24 is a synthetic strigolactone analog commonly used in laboratory germination studies of parasitic plants as a positive control. 5-deoxystrigol is one of the major SLs produced by compatible grass hosts (Awad et al., 2006) and is a potent stimulator of parasite germination whereas orobanchol is a more dominant SL among dicot hosts (Yoneyama et al., 2008) and is a less potent stimulator of *S. hermonthica* germination for certain genotypes (Haussmann et al., 2004; Cardoso et al., 2014; Bellis et al., 2020).

In addition to seed collections kept separately from individual plants, we also included tests of bulk seed collected from the Kibos population that were confirmed to have high germinability in our previous experiments (Bellis et al., 2020). We used a generalized linear mixed model (GLMM) with a random effect of *S. hermonthica* genotype (of the parent plant) to compare differences in germination rate among sites, hosts, and treatments (orobanchol vs. 5- deoxystrigol). GLMMs were implemented in R version 4.0 with the lme4 package (Bates *et al*. 2015), and significance of fixed effects was determined using likelihood ratio tests.

## RESULTS

### Sequencing

For the 68 Kenyan samples, on average 80% of reads per library were classified as plant-derived using Kraken 2 (range: 71-86%). Most classified reads were assigned specifically to *S. hermonthica* (mean: 46%) or *S. asiatica* (mean: 30%), both represented in our database by high-quality transcriptomic or genomic resources from which contaminant sequences (including from plant hosts) have been carefully filtered (Yang et al., 2014; Yoshida et al., 2019). After quality and adapter trimming, on average 7.7 Gigabase pairs of plant-derived sequence data remained per sample (range: 5.2-11.7 Gbp). Given flow cytometry estimates of the genome size of *S. hermonthica* ranging from 1C = 0.9 Gb (Yoshida et al., 2010) to 1C = 1.4 Gb (Estep et al., 2012), this sequencing effort corresponds to an approximate average depth of 5.5x to 8.6x read coverage per base for each sample.

### Population structure

Principal Coordinates Analysis (PCoA) based on *k*-mers indicated high genetic diversity within populations, with only subtle differentiation of populations from the same farm collected from different hosts (Fig. 1). Greater correlation was observed between genetic and geographic distance than expected by chance (*p* = 0.001, Mantel test), but patterns of isolation-by-distance were extremely weak (Fig. 1F). Within-population diversity was very high at the geographic scale investigated. Many individuals exhibited a comparable range of pairwise genetic diversity within a single field (*k*-mer distance ranging from 0.0216 - 0.0286; *n* = 572 pairwise comparisons) as between individuals sampled more than 100 km apart (*k*-mer distance ranging from 0.0242 - 0.0286; *n* = 336 pairwise comparisons; Fig. 1F).

Our alignment-free approach may not remove all contaminant sequences, and so we also carried out a reference-based analysis using a draft reference genome assembled as part of this study. Estimation of pairwise distances from reference-based analyses suggested that samples were not strongly clustered by host or by sampling location, except for nine individuals parasitizing finger millet in Kisii that clustered separately from other samples (Fig. 2). These findings are consistent with our *k*-mer based analysis, in which the same individuals appear close together and are also separated from other finger millet parasitizing individuals along the first PCoA axis (Fig. 1D). Although some limited population genetic structuring may be present within highly diverse *S. hermonthica* populations (perhaps due to multiple source populations, for example), populations are not strongly differentiated by host at the genome-wide level.

**Figure 2.**
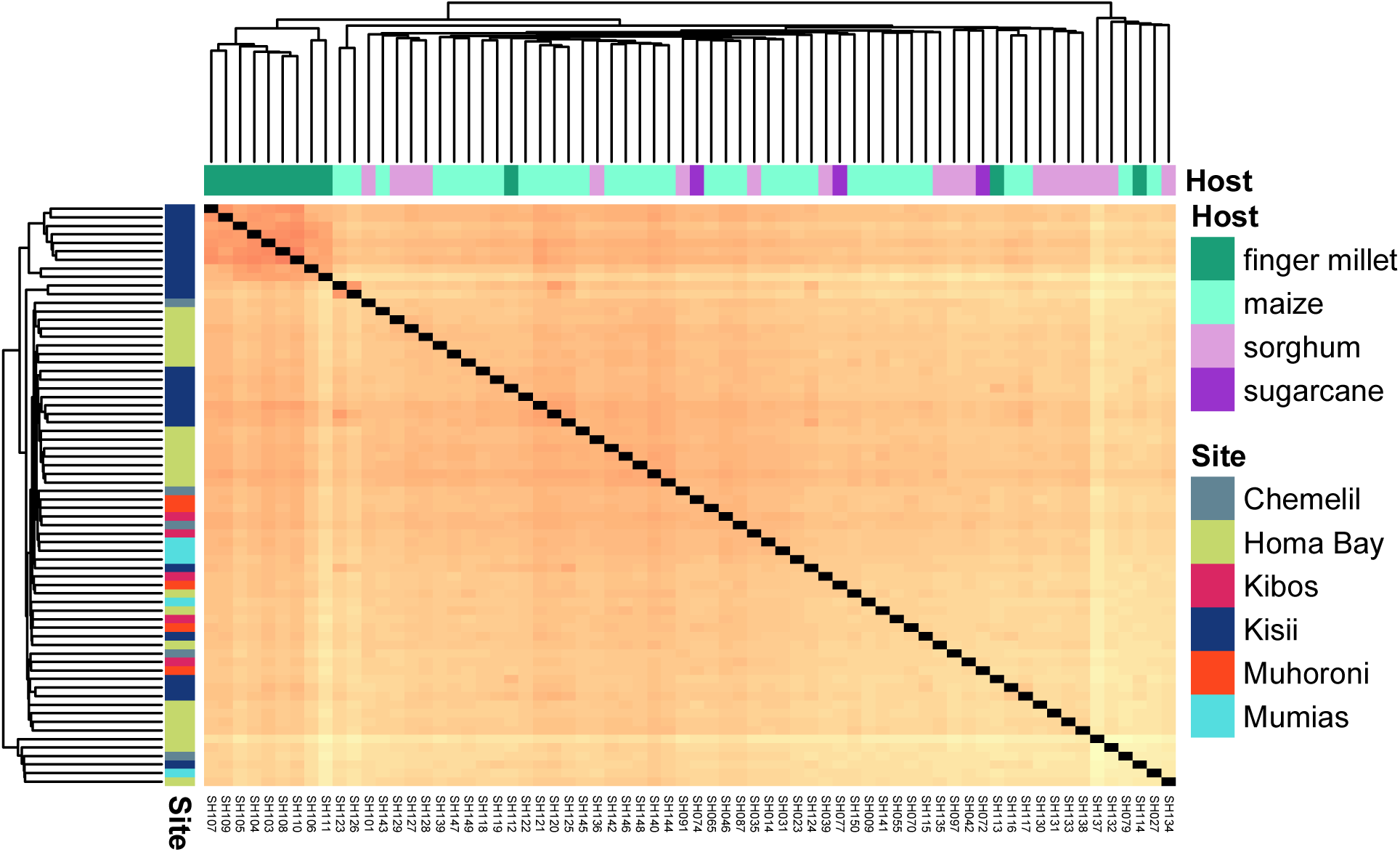
Population structure of Kenyan *S. hermonthica* samples from reference-based analysis. Pairwise genetic distances were estimated from genotype probabilities, for 11,615,822 variable sites across 43,910 contigs longer than 2.5 kb. Genetic distances are colored in a heat map-scale from 0.24 (red) to 0.33 (yellow). Individuals are ordered based on a hierarchical clustering computed from the estimated genetic distances.

### Germination rate variation

Previous studies have suggested that in contrast to other locations in Africa, *S. hermonthica* populations from western Kenya demonstrate a generalist germination response to strigolactones (Haussmann et al., 2004; Bellis et al., 2020). However, *Striga* germination tests are typically conducted with bulk seeds collected from many individuals in a field, potentially masking individual-level variation that could be segregating with respect to parasitism on different hosts, for example if a generalist population is composed of individuals that specialize on different resources (Bolnick et al., 2002).

To characterize individual-level variation in *S. hermonthica* parasitism, we conducted controlled germination tests in the USDA-permitted quarantine facility at Pennsylvania State University. Positive controls with bulk seed (collected in the Kibos region) and 0.2 µM of the artificial strigolactone GR24 indicated good germinability for positive controls (66.5%), and no germinated seeds were observed in wells with only sterile water. Compared to the higher concentration of GR24, bulk seed showed slightly lower germination rates in response to 0.01 µM of the natural SLs orobanchol (51.0%) and 5-deoxystrigol (55.7%) indicating that the concentrations of orobanchol and 5-deoxystrigol used in the individual-level experiment should produce strong but sub-maximal germination responses.

We did not find significant host-associated germination variation among seeds collected from individual parasites on different hosts (Table S1; Fig. 3). After accounting for treatment and site, host-of-origin was not a statistically significant effect in our model (*P* = 0.247; likelihood ratio test for test of full vs. reduced GLMM). The probability of germination was 30% lower for individuals from Kisii compared to Homa Bay (*P* = 0.04; GLMM with fixed effects of ‘Site’ and ‘Treatment’) and 15% lower in response to orobanchol than to 5-deoxystrigol (*P* = 0.004; GLMM).

**Figure 3.**
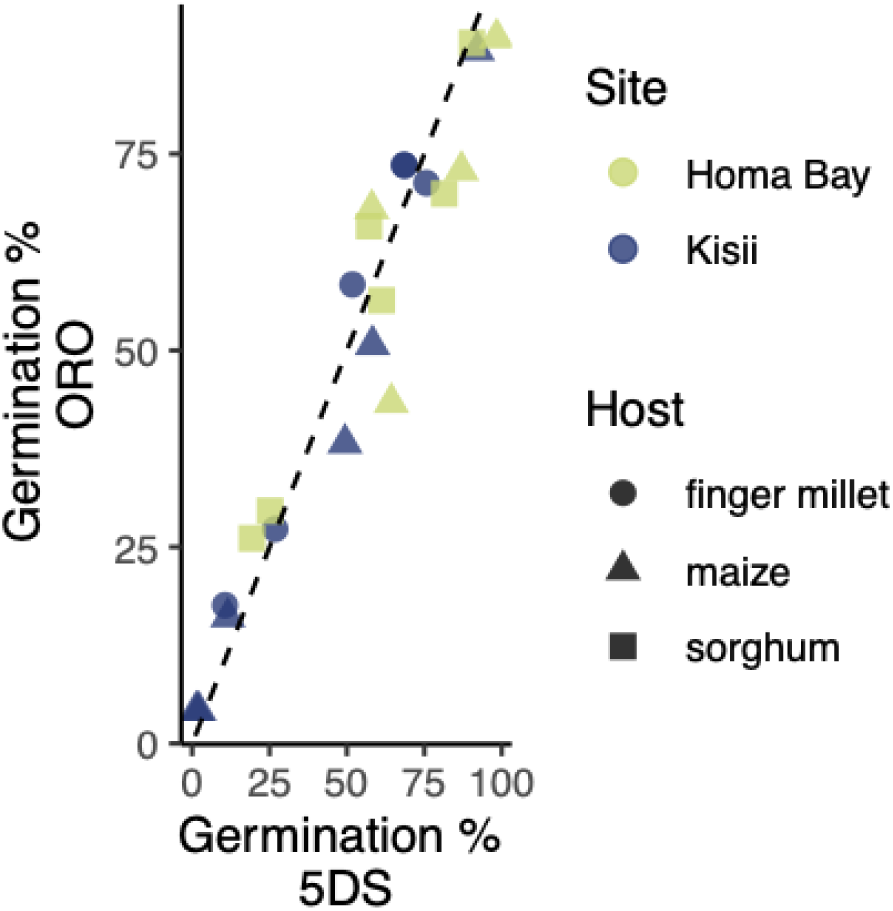
Germination response variation in Homa Bay and Kisii populations. Seeds from *n* = 6 individuals per unique site and host were tested in response to synthetic strigolactones orobanchol (ORO, 0.01 μM) and 5-deoxystrigol (5DS, 0.01 μM). The dashed line indicates the expectation if the percent germination in response to the two different germination stimulants is identical.

### Host-associated loci

Germination tests did not support host-specific differences in strigolactone response in these populations. However, it is possible that host-specific selection could still leave detectable signatures at the genetic level. We identified host-associated genetic variation without a reference using a *k*-mer based approach. For Homa Bay, the total number of *31*-mers generated was 2,765,562. For Kisii, the total number of *31-*mers generated was 2,123,902. Highly differentiated *k*-mers were defined as those having *G_ST_* > 0.5. A lower proportion of *31*-mers were highly differentiated for parasites on sorghum vs. maize hosts in Homa Bay (0.6% of *31*-mers) compared to finger millet vs. maize hosts in Kisii (4.8% of *31*- mers; Fig. 4A). These proportions far exceeded the proportions expected by chance, based on *G_ST_* values observed among 1000 random permutations of host labels for each of the populations (Fig. 4B, D). In 1,000 random permutations, the maximum proportion of *k*-mers with *G_ST_* > 0.5 for the Homa Bay population was 0.2%, whereas the maximum for the Kisii population was 0.5%.

**Figure 4.**
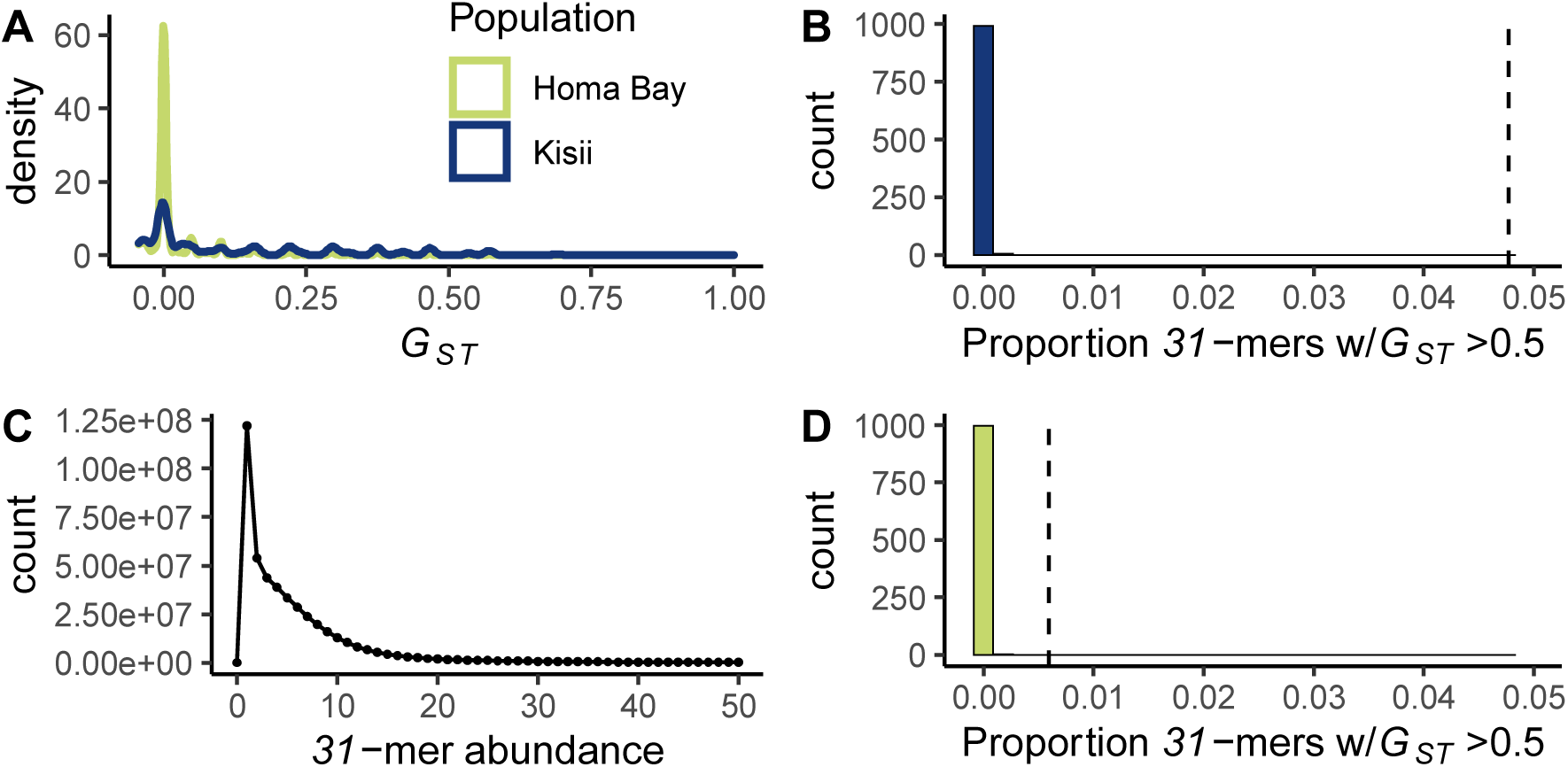
*G_ST_* and *k-*mer distributions. A) Observed distribution of *G_ST_* values. For Homa Bay, *G_ST_*is for parasites in adjacent plots of maize vs. sorghum at 2,765,562 *31*-mers. For Kisii, *G_ST_* is for parasites in adjacent plots of maize vs. sorghum at 2,123,902 *31*-mers. B) Proportion of *31-*mers with *G_ST_* >0.5 in the Kisii dataset. The shown distribution is based on 1,000 random permutations of host labels. The dashed line indicates the observed proportion. C) A representative *k-*mer distribution spectrum for sample SH046. Most unique *31*-mers are observed only once or twice due to the high heterozygosity of this species. D) Proportion of *31-* mers with *G_ST_* >0.5 in the Homa Bay dataset. The shown distribution is based on 1,000 random permutations of host labels. The dashed line indicates the observed proportion.

Highly differentiated *31-*mers were then assembled into longer contigs for follow-up analysis. For the Homa Bay population (maize vs. sorghum hosts), 42 contigs were assembled ranging in length from 58 to 91 bp. Fifteen of these contigs had good BLAST hits to the transcriptome database, with >85% identity over at least 45 bp (Table S2). For the Kisii population (finger millet vs. maize hosts), highly differentiated *k*-mers assembled into 469 contigs with length ranging from 29 to 211 bp. Of these, 240 had good BLAST hits to transcripts in our database (Table S3).

Of particular interest to our investigation were host-specific contigs with high levels of similarity to known *Striga* parasitism genes including SL receptors and genes involved in haustorium development. We first searched for similarity to a set of 11 *S. hermonthica* strigolactone receptors that vary in their binding affinity for diverse strigolactones [GenBank accession numbers KR013121 – KR13131] (Tsuchiya et al., 2015). We observed only one likely spurious match to *ShHTL2* (87% similarity over 23 bp) and no hits to any of the 21 *KAI2* paralogs from the *Striga asiatica genome* (Yoshida 2019). In follow-up analyses based on alignments to reference *ShHTL* transcripts, only one site in *ShHTL6* had an estimated difference in allele frequency between parasites on finger millet and maize exceeding 0.5 (Fig. S1). This polymorphism occurred at position 457 and does not result in an amino acid change. Together, we find little phenotypic or genomic evidence supporting host-specific differentiation for loci involved in perception of strigolactones in these populations.

In contrast, several assembled host-specific contigs had good matches to loci implicated in development of haustoria. Annotated transcripts included a 60 bp contig with >98% similarity over its full length to a transcript annotated as *SUPPRESSOR OF G2 ALLELE SKP1* (*SGT1*) (Table S2). *SGT1* was among the top upregulated genes in haustoria of the root parasitic plant *Thesium chinense* and was hypothesized to be important for generating auxin response maxima during haustorium development (Ichihashi et al., 2017). In *S. hermonthica,* it is also highly expressed in imbibed seeds (Stage 0) and in haustoria attached to host roots (Stage 3; Fig. S2).

Parasitism on different hosts was associated with genetic structural variation in *SGT1*, with two <100 bp regions often absent from parasites on sorghum but present for parasites on maize and finger millet. We also identified a 59 bp contig with 96% similarity over its full length to a transcript annotated as a pectin methylesterase; pectin methylesterases have previously well-characterized functions in developing haustoria (Yang et al., 2014). The pectin methylesterase transcript was not expressed in parasites grown on sorghum in most stages, but exhibited low, non-zero expression in *S. hermonthica* seedlings after exposure to a haustorium inducing factor (mean TPM = 0.03). Strong differentiation in transcript-aligned reads between individuals from finger millet (frequency of the deletion allele = 0.65) and maize (frequency of the deletion allele = 0.03) verified the signal observed from the *k*-mer association analysis.

Two additional assembled host-differentiated contigs of 113 bp and 103 bp had 99% and 97% similarity, respectively, over their full length to separate regions of a single transcript annotated as a precursor of chemocyanin (Table S3). Chemocyanins may be of particular interest due to the evolutionary co-option of many pollen tube genes by parasitic plants for haustorium development (Yang et al., 2014) and the key role of chemocyanin as an attractant for directing pollen tube growth (Kim et al., 2003). Alignment of our DNA sequences to the chemocyanin transcript reference revealed host-associated structural variation (Fig. 5A). PCR-based confirmation indicated 500 bp or more of genome sequence directly upstream of the 5’ end of the transcript was completely missing from parasites on maize, suggesting potential impact on variation in gene expression levels. The fragmented nature of the draft genome assembly precluded our ability to design PCR primers completely spanning the deletion, but PCR banding patterns showed a characteristic absence of the region in individuals having the allele more common on finger millet (Fig. S3), and a diversity of deletion alleles at this site (Fig. S4). Using just sequences for individuals parasitizing finger millet, we reconstructed a 934 bp contig, from reads that mapped to the transcript reference and their unmapped pairs.

**Figure 5.**
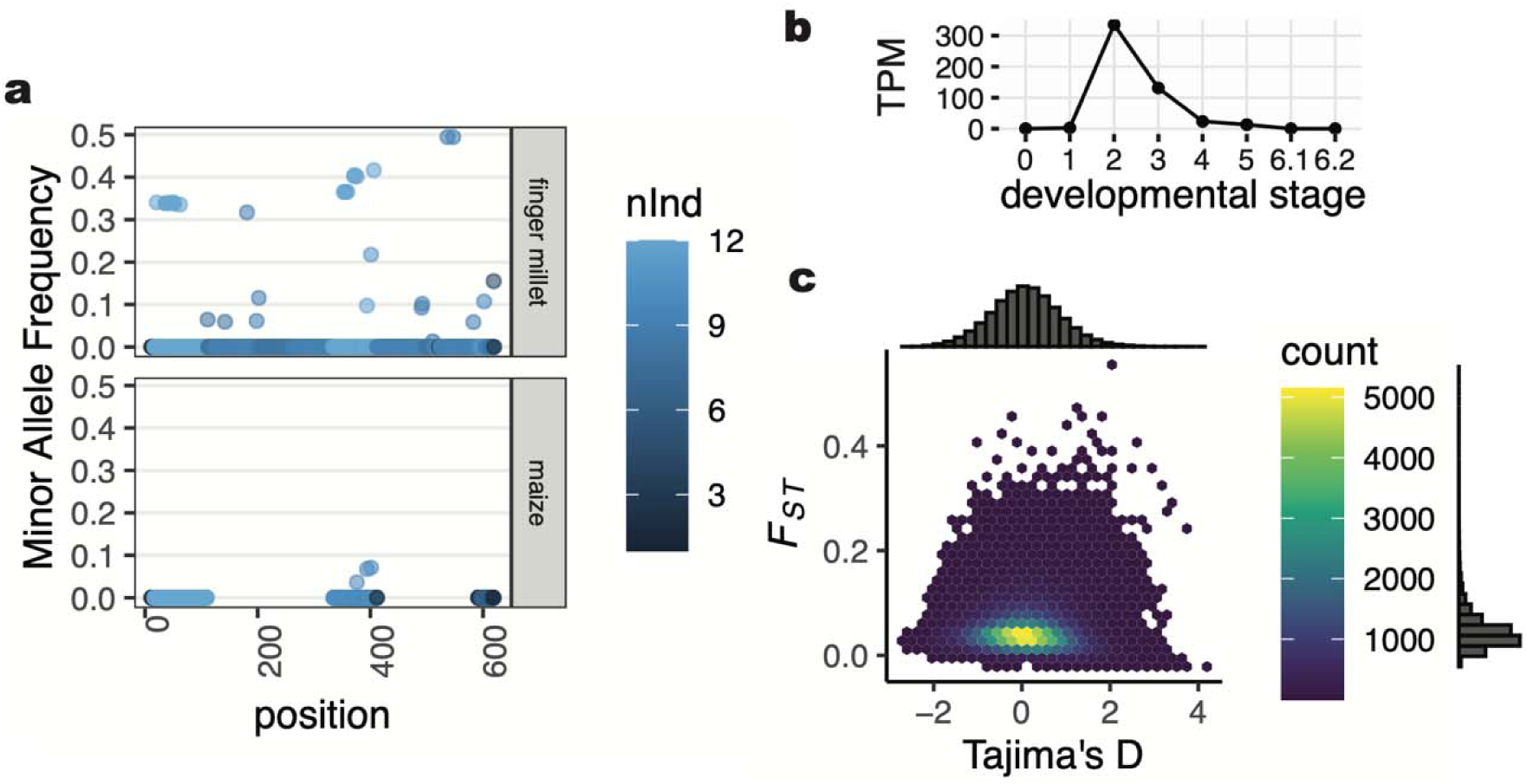
Genetic variation in *Striga hermonthica* from Kisii, Kenya. (a) Presence/absence variation across the chemocyanin precursor transcript (StHeBC4_h_c11261_g0_i1), for *n* = 12 individuals per host (finger millet or maize). If the locus is not present in any of the sequenced individuals, no data point is shown, otherwise each point is colored according to the number of individuals (nInd) with data for the position. Two regions in the transcript have no reads aligned from maize-parasitizing individuals (lower panel), consistent with absence of the corresponding *k*-mers and PCR-banding results that suggest a deletion. (b) Gene expression data in transcripts per million (TPM) from the PPGPII dataset for the chemocyanin precursor transcript across six stages of haustorial development (0: imbibed seed; 1: germinated seedling after exposure to GR24; 2: germinated seedling after exposure to DMBQ; 3: ∼48 hrs post-attachment; 4: ∼72 hrs post-attachment; 5: late post-attachment; 6.1: vegetative structures; 6.2: reproductive structures). (c) Distribution of Tajima’s D and *F_ST_* values across 154,722 non-overlapping 1-kb windows. Only windows with data for more than 50% of sites are shown, excluding impacts of structural genetic variation as in (a). Tajima’s D for the chemocyanin was - 1.7.

Alignments to this ‘finger millet’ allele confirmed its presence in 9/12 individuals from Kisii parasitizing finger millet and 0/12 parasites on maize (Table S3). Among all individuals sequenced in our study, the allele is present at lowest frequency in maize parasites (14% of individuals; *n* = 35) and intermediate frequency for parasites on sorghum (22%; *n* = 18); and sugarcane (33%; *n* = 3)

Although patterns of allele frequency differentiation are partially correlated with overall patterns of population structure (Fig. 2), additional lines of evidence point to an important role of the chemocyanin in parasitic plant-host interactions. The chemocyanin transcript was present in a previously identified list of *S. hermonthica* “core parasitism” genes, with highest expression after exposure to haustorial initiation factors and during early post-attachment (Yang *et al*. 2014; Fig 3B). It was also highly expressed in cells at the host-parasite interface in a study that used laser capture microdissection to characterize gene expression in the *S. hermonthica-*sorghum interaction (Honaas et al., 2013).

### Validation of k-mer-based approaches

To further investigate signatures of selection in the context of genome-wide patterns, we assembled a draft genome for *S. hermonthica* from South Uganda. The total length of the assembled genome after filtering was 1,431 Mbp over 12,155,247 contigs, with a maximum contig length of 37.5 kb. The assembly was highly fragmented with the largest 1.69 million contigs accounting for 50% of the assembly, and the length of these contigs ≥ 110 bp (N/L50 = 1690935/110). Nevertheless, sequences of interest were present on contigs long enough to allow for further interrogation. Specifically, the assembled chemocyanin transcript for the finger millet allele had 97.8% identity over 918 bp to a single contig of 1,055 bp (NODE_132909_length_1055_cov_32.632), and no other close hits.

After removing contigs shorter than 500 bp, 307026 scaffolds with a total length of 479 Mbp remained.

To evaluate signatures of selection on the chemocyanin gene in the context of genome-wide patterns, we estimated Tajima’s D and *F_ST_* for all assembled 1-kb non-overlapping windows for maize- vs. finger millet-parasitizing individuals in the Kisii population. Consistent with the expected signature of a selective sweep, a Tajima’s D value of -1.7 in the Kisii population indicated a significant excess of low-frequency polymorphism (empirical *P*-value = 0.009) for the chemocyanin contig in the *de novo* assembly compared to the genome-wide value of 0.11 ± 0.013 (mean ± std. error; linear mixed effects model; distribution of Tajima’s D values relative to *F_ST_* shown in Fig. 5C). For non-overlapping 1-kb windows with data (*n =* 237,632 windows), the median *F_ST_* value was 0.038. Of these windows, 1.8% had an average *F_ST_* value greater than 0.15 and 0.5% had an average *F_ST_* value greater than 0.2. The maximum value of *F_ST_* observed for the reference-based analysis was 0.57, for a 4-kb contig with similarity to a *Phelipanche aegyptiaca* gene annotated as an ortholog of AT5G64440 (fatty acid amide hydrolase) which is an important regulator of plant growth (Wang et al., 2006). Notably, *F_ST_*for chemocyanin contig could not be reliably calculated using the reference-based approach because only 30 positions in the 1055-bp contig had data for a majority of individuals due to structural variation. Together with observations from the *G_ST_* analysis, our results suggest that only a small portion of the *S. hermonthica* genome is strongly differentiated by host species in these populations from western Kenya.

## DISCUSSION

Agricultural weeds are increasingly recognized as important model systems for addressing key questions in evolution and ecology (Vigueira et al., 2013; Baucom, 2019). In particular, compared to many other systems, parasitic weeds offer particular advantages for the study of coevolution including well-developed genomic and germplasm resources for their hosts, high quality distributional data, and a rich literature describing variation in species interactions over many decades (Bellis et al., 2020, 2021). Yet, despite recent advances in genome sequencing for parasitic plants, evolutionary analyses particularly for species with large, complex genomes (e.g. >1 Gb) remain a challenge (Lyko and Wicke, 2021). Consequently, previous population-level diversity studies for *S. hermonthica*, one of the most damaging parasitic plants in agriculture, have used reduced representation approaches (Unachukwu et al., 2017; Lopez et al., 2019). However, reduced representation approaches such as RAD-seq may miss signatures of selection that are highly localized in the genome (Lowry et al., 2017; Lou et al., 2021), and transcriptomes fail to provide information regarding non-coding regions of the genome, which also generate phenotypic diversity. As the cost of sequencing continues to decrease, whole genome resequencing coupled with alignment-free bioinformatic approaches can provide a promising alternate approach for surveying population genomic diversity (Voichek and Weigel, 2020). Here, we report some of the first publicly available WGS data for field-sampled individuals of the parasitic weed *Striga hermonthica*. Our analyses underscore high within-population genomic variation and implicate host-specific selection on genes involved during parasite attachment and haustorial development.

Perhaps surprisingly, we find little genomic or phenotypic evidence for host-specific selection on strigolactone perception variation in the studied populations. Specifically, one may expect selection on perception loci to be relatively strong, particularly since *S. hermonthica* is an obligate parasite and so the costs of germination in the absence of a suitable host are high. This suggests that, at least in western Kenya, most common grass hosts may be suitable, so that costs of germinating on the wrong host are low. The genomes of *Striga* spp. include a diverse repertoire of strigolactone receptors, each with variable affinity for different SLs (Tsuchiya et al., 2015; Yoshida et al., 2019), providing many potential targets for selection on SL response variation. One possibility is that these receptors are now subject to purifying selection in western Kenya, rather than diversifying selection expected if SL perception variation is strongly linked to fitness variation across different hosts. Environmental niche models from our previous study (Bellis et al., 2021) predict highest suitability for *Striga* parasitism of maize across sampling locations in our study (mean suitability: 0.92) but lower suitability for sorghum and pearl millet (0.46 and 0.03, respectively). Another possibility is that the particular host genotypes studied here may overlap in SL exudation profile enough that selection on parasite germination rate variation is not strong in these natural field populations. For example, while zealactones appear to be uniquely produced by maize (Charnikhova et al., 2017), some varieties also naturally produce sorgomol and 5-deoxystrigol in high quantities (Yoneyama et al., 2015), strigolactones common in sorghum root exudate that promote strong germination response in *S. hermonthica*.

A third explanation, if selection on SL perception is indeed important for local adaptation, is that selection may only be strong in some populations across the range of *S. hermonthica*. This explanation is consistent with the idea of coevolutionary hotspots, where there is reciprocal selection among coevolving species (Thompson, 2005), but with the genetic targets of selection involving different stages of the infection process in different locations. In western Kenya, for example, coevolutionary hotspots may be ‘hotter’ for genes involved in parasitic interactions post- germination than for SL perception. Kenyan *S. hermonthica* populations exhibit a more generalist germination response compared to populations from West Africa, which show pronounced differences in response to orobanchol vs. 5-deoxystrigol from host root exudates or to strigolactone standards (Parker and Reid, 1979; Haussmann et al., 2004; Bellis et al., 2020). This idea also corresponds with previous findings that East African *S. hermonthica* may have greater average infestation success across diverse sorghum genotypes than West African populations (Bozkurt et al., 2015). Functional G_H_ x G_P_ in strigolactone response variation may segregate among populations in a different part of the parasite range than studied here, for example among West African populations, or at a broader spatial scale, for example in West vs. East African parasite populations (Haussmann et al., 2004; Bellis et al., 2020).

In contrast to general expectations (Nuismer and Dybdahl, 2016), our genomic analyses revealed the strongest evidence for host-specific selection on genes involved in the later stages of parasite development. The best evidence for host-specific selection on haustorium loci came from our analyses of a transcript annotated as a chemocyanin precursor (Fig. 5). In addition to strong differentiation between finger millet and maize hosts, the 1-kb region including the chemocyanin exhibited an excess of rare polymorphism and multiple alleles (Fig. S3-S4), consistent with expectations for recurrent soft sweeps from standing genetic variation (Pennings and Hermisson, 2006). Notably, even the strongest signals of host-specific selection detected in our study did not reveal any loci exhibiting complete or near complete differentiation between parasites on different hosts, indicating that there may be relatively few genetic barriers to parasitism of different cereal host species in the studied region. The ‘finger millet’ chemocyanin allele, for example, was also present at low frequency in the genomes of parasites on other host species. This suggests a neutral impact of the allele on parasitism of other hosts, and potentially, a lack of trade-offs, given that all sequenced parasite individuals were already at an advanced life stage. The importance of conditional neutrality for local adaptation in other systems (Lowry et al., 2019) further highlights the complexity of selective pressures shaping local adaptation of parasitic plants to dynamic host communities.

## CONCLUSIONS

While our results emphasize the challenges of *Striga* management due to high genomic diversity and adaptive potential, they also highlight the promise of low coverage WGS approaches for functional genomics of non-model species. The outlier signal for two of the three candidate loci we describe in detail here resulted from structural variation that would not have been uncovered in an alignment-based analysis or using a RAD-Seq approach that may only survey a small portion of the genome or be prone to allele drop-out. Reference-free approaches continue to gain ground for studies of genomic and phenotypic variation in plants, with well-documented advantages (VanWallendael and Alvarez, 2020; Voichek and Weigel, 2020). Our study indicates that the utility of large WGS datasets may not be out of reach even for species such as *Striga hermonthica* characterized by large, complex genomes. As the need to mitigate biotic constraints on global food security becomes increasingly critical, reference-free analyses coupled with WGS data may serve as a promising strategy for rapid characterization of alleles involved in parasite adaptation across diverse environments.

## ACKNOWLEDGMENTS

This study is based on work supported by a National Science Foundation Postdoctoral Research Fellowship in Biology to E.S.B. under Grant 1711950, the Arkansas Biosciences Institute (the major research component of the Arkansas Tobacco Settlement Proceeds Act of 2000), and the *Emmy Noether*-program of the German Science Foundation (WI4507/3-1 to S.W.). A.K. and X.H. were supported by NSF EPSCoR Award 1946391. J.R.L. was supported by NIH R35 GM138300. This work is supported by the Arkansas High Performance Computing Center which is funded through multiple National Science Foundation grants and the Arkansas Economic Development Commission. We also thank the Texas A&M AgriLife Research: Genomics and Bioinformatics Service for library preparation and sequencing.

## AUTHOR CONTRIBUTIONS

C.S.v.M., S.W., C.O.O., E.K., T.X., and E.S.B collected and processed samples and performed laboratory experiments. C.S.v.M., S.W., A.K., and E.S.B. carried out analyses. X.H., S.W., S.M.R., C.W.D., J.R.L., and E.S.B. contributed to experimental design and interpretation. E.S.B. and J.R.L. drafted the manuscript, with input and critical revision from all authors. All authors approved the final version of the manuscript.

## DATA AND CODE AVAILABILITY

Raw reads from whole genome sequencing of the 68 *S. hermonthica* individuals from the 2018 collection and the Ugandan reference have been deposited in the National Center for Biotechnology Information (NCBI) Sequence Read Archive (SRA) database, https://www.ncbi.nlm.nih.gov/sra (BioProject accession no. PRJNA801489). The reference assembly of the Ugandan specimen is available for download and BLAST searches on *WARPP* (https://warpp.app; (Kösters et al., 2021)). Images associated with the different collection sites are available from iNaturalist. Germination rate data and code to reproduce the analyses are available at https://github.com/em-bellis/StrigaWGS.

## SUPPLEMENTARY INFORMATION

Additional supporting information may be found online in the Supporting Information Section at the end of the article.

Appendix S1: Supplementary Figures and Tables.

## Appendix S1: Supplementary Figures and Tables

**Figure S1.**
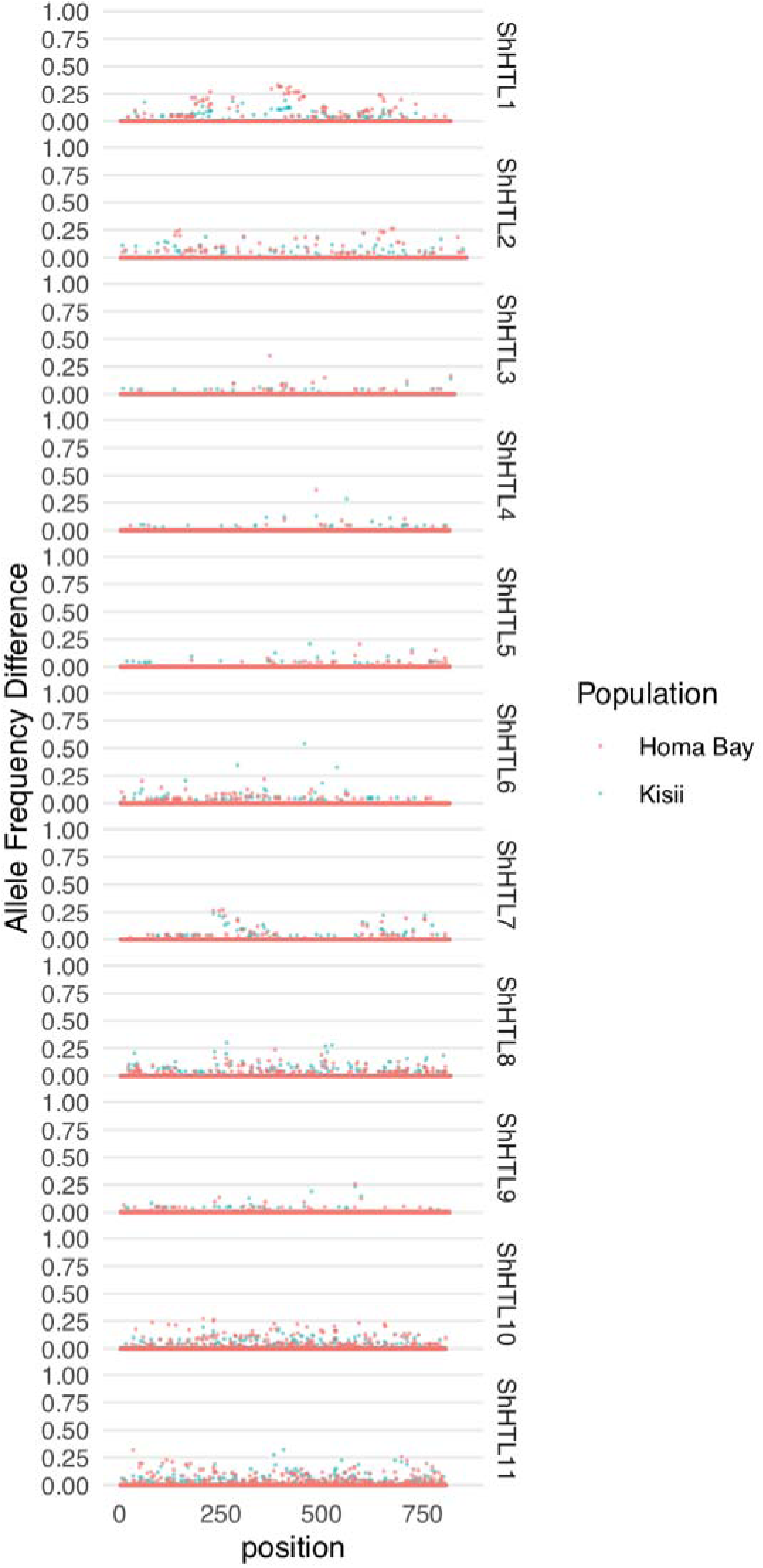
Allele frequency differences for 11 *ShHTL* receptors from Tsuchiya et al. (2015). Allele frequencies were estimated from genotype likelihoods based on reads mapped to each reference transcript. Allele frequency was estimated for parasites on each host species separately (*n* = 12 per unique host and population). The difference in allele frequency estimates between two hosts within a single population is shown.

**Figure S2.**
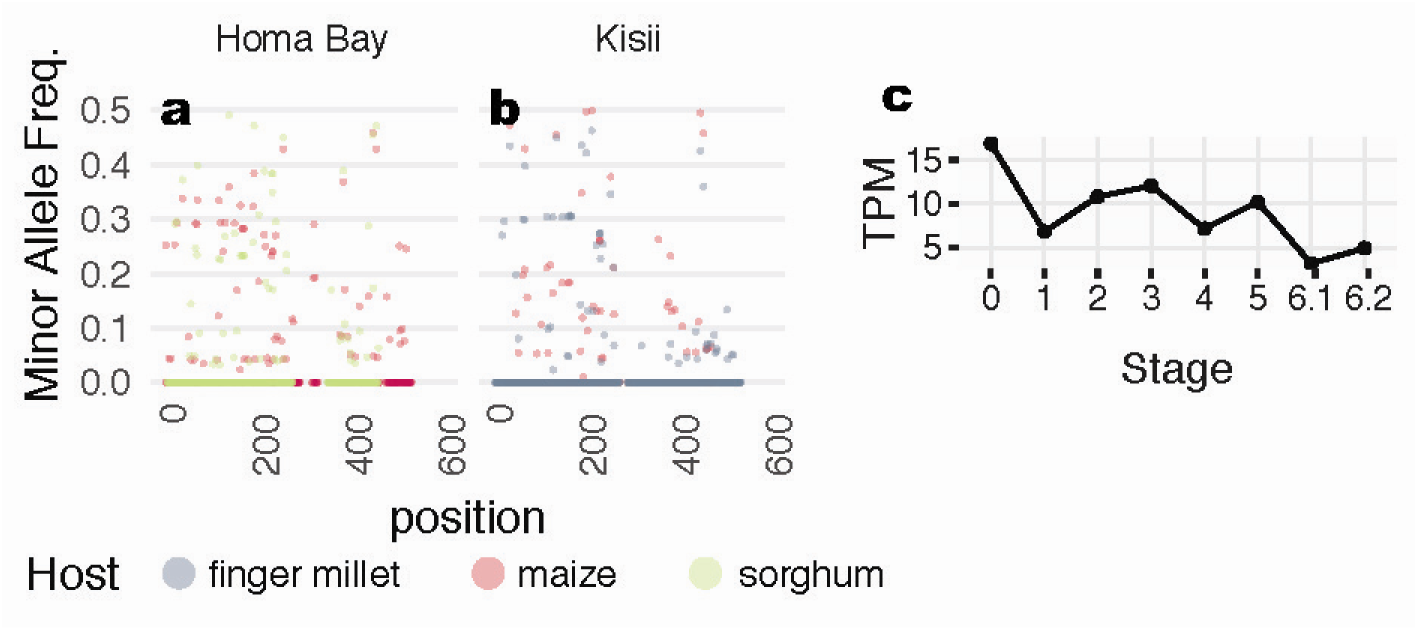
(a, b) Minor allele frequency differences based on alignments to the *SGT1* transcript sequence (StHeBC4_p_c12587_g2_i1), split by population. If the locus is not present in any of the sequenced individuals for that population due to genetic structural variation, no data point is shown. Frequencies were estimated based on genotype likelihoods from *n* = 12 individuals per unique host and population. (c) Gene expression data in transcripts per million (TPM) from the PPGPII data across the 6 stages of haustorial development from data published.

**Figure S3.**
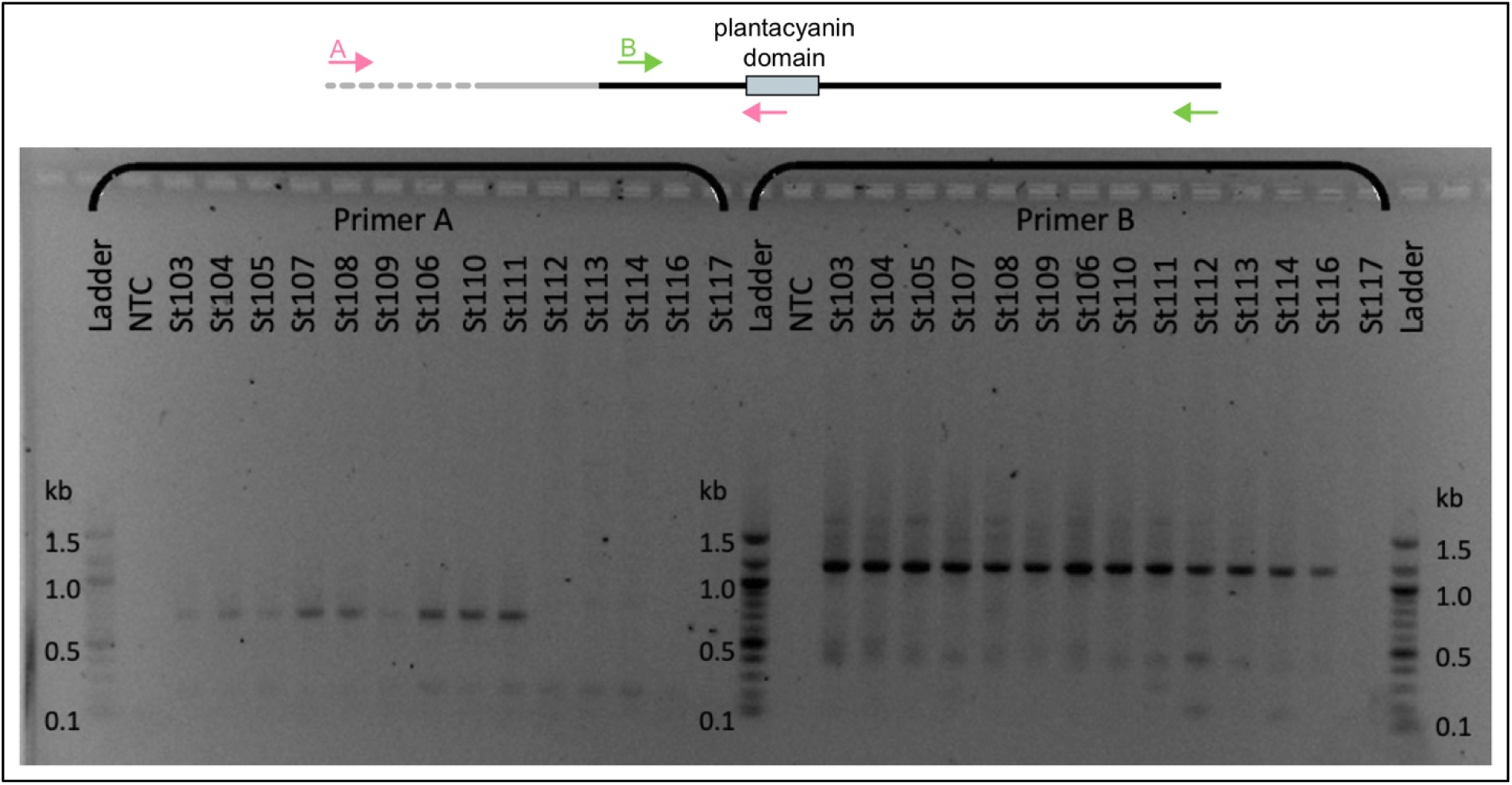
PCR banding results to confirm chemocyanin deletion calls. In the top panel, binding sites for primer sets A (pink) and B (green) are shown relative to location of the transcript sequence (solid black line), location of the deletion (grey dashed line), and conserved plantacyanin domain (light blue rectangle). Primer set A does not yield a 676-bp band for individuals with the deletion (lanes 11-15) whereas primer set B amplifies a region downstream of the deletion. NTC: no template control.

**Figure S4.**
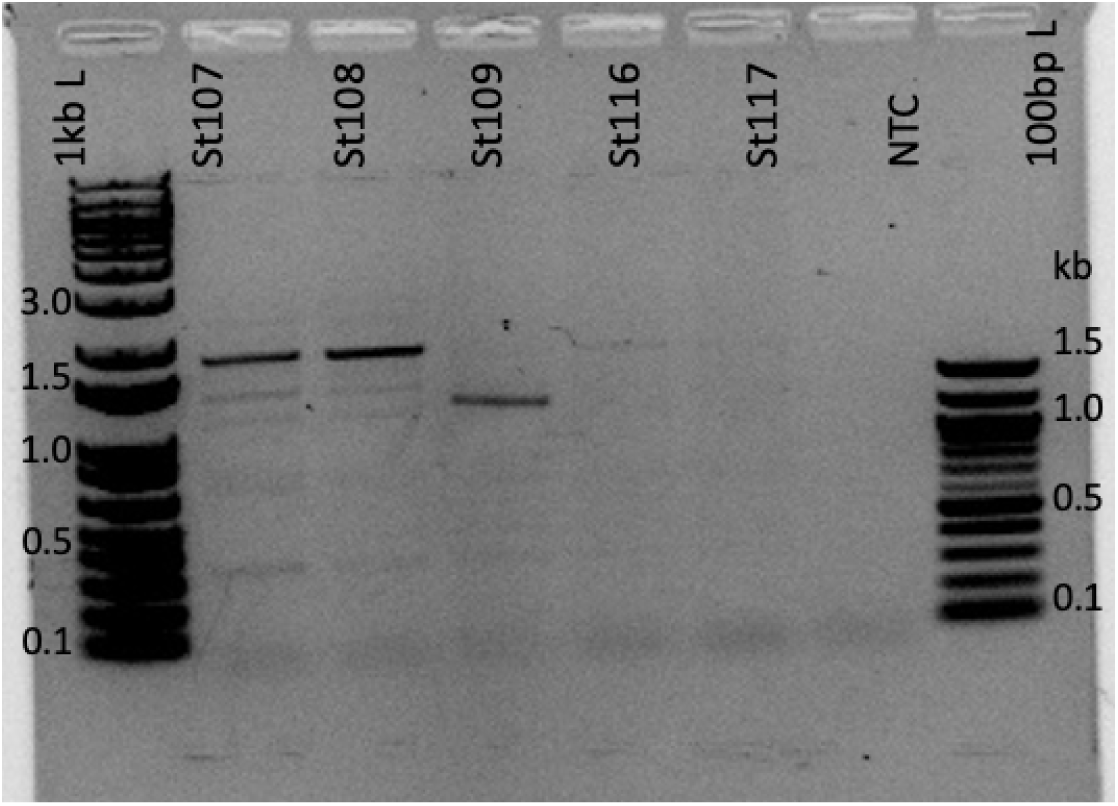
PCR banding results suggest multiple deletion variants. PCR amplification using the outer primers from set A (5’-TCTCGATCCTTTTGGAATGG-3’) and B (5’- TGGGGAAAGAGGTAGTGCAA-3’); see Fig. S3). NTC: no template control. Lane 1: Sh107; Lane 2: Sh108, Lane 3: Sh109; Lane 4: Sh116; Lane 5: Sh117.

**Table S1.** GLMM for analysis of germination rate variation

|  | **Estimate** | **Std. Error** | **z-value** | ***P*** |
| --- | --- | --- | --- | --- |
| Intercept | 2.29 | 0.97 | 2.37 | 0.017 |
| Site == Kisii | -2.16 | 0.79 | -2.72 | 0.006 |
| Host == Maize | -1.13 | 0.79 | -1.43 | 0.153 |
| Host == Sorghum | -1.90 | 1.12 | -1.70 | 0.089 |
| Treatment == ORO | -0.16 | 0.06 | -2.85 | 0.004 |

**Table S2.**
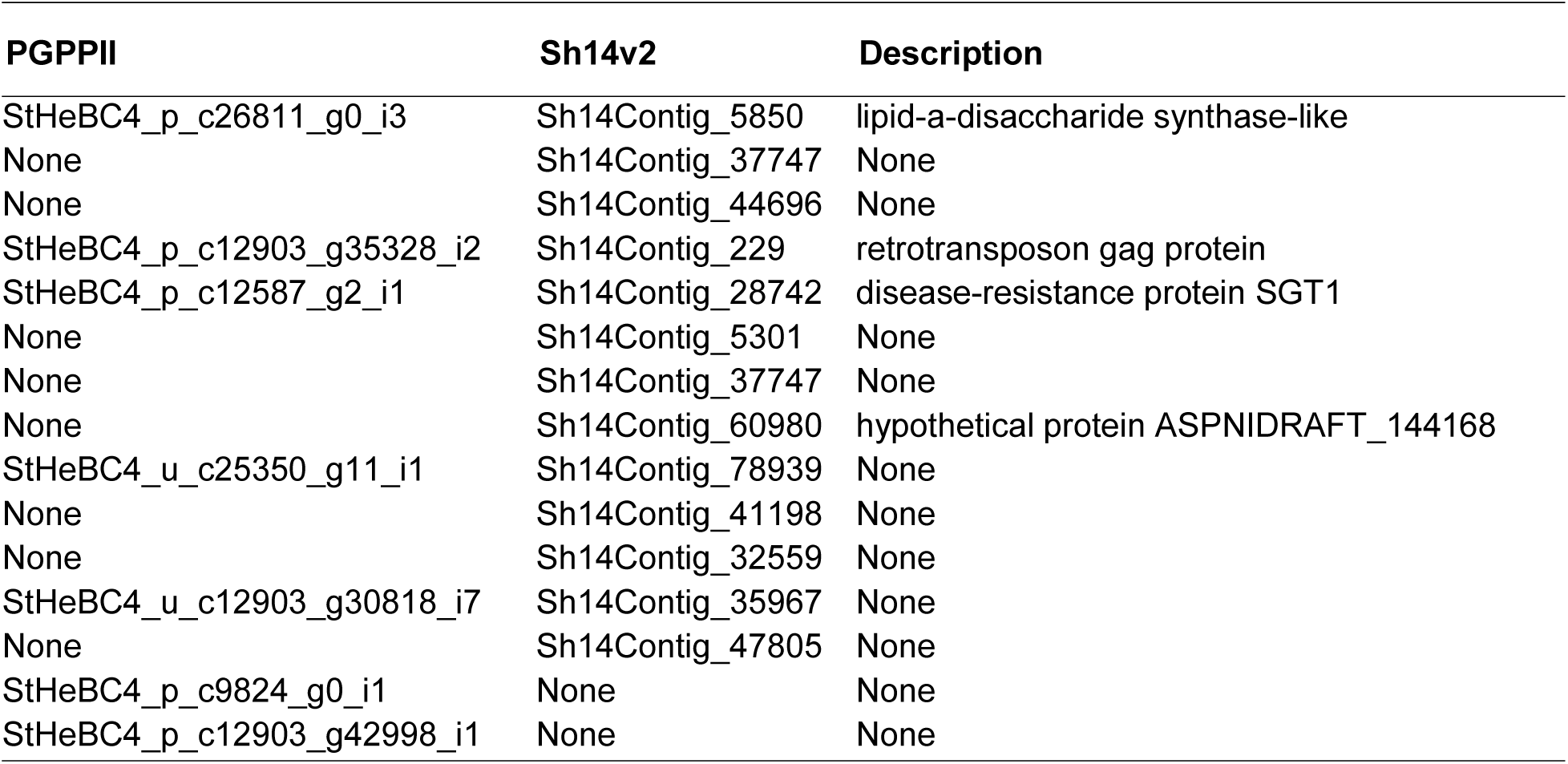
Hits to transcripts from the Parasitic Plant Genome Project (PPGPII) or Sh14v2 (from Yoshida 2019) for contigs assembled from host-associated *k-*mers for *Striga hermonthica* Homa Bay population (maize vs. sorghum hosts). Transcripts with greater than 85% identity over at least 45 bp are shown.

**Table S3.**
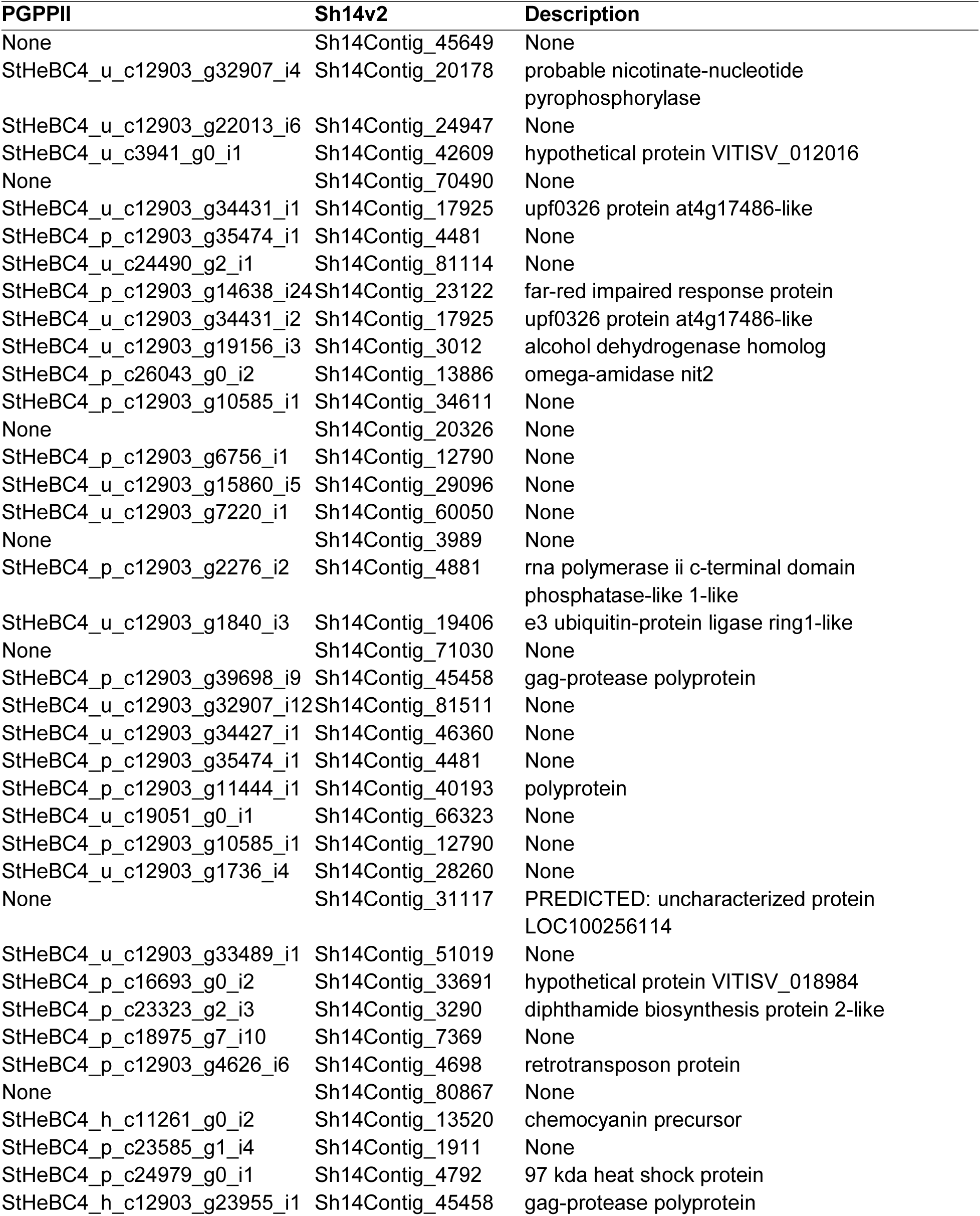
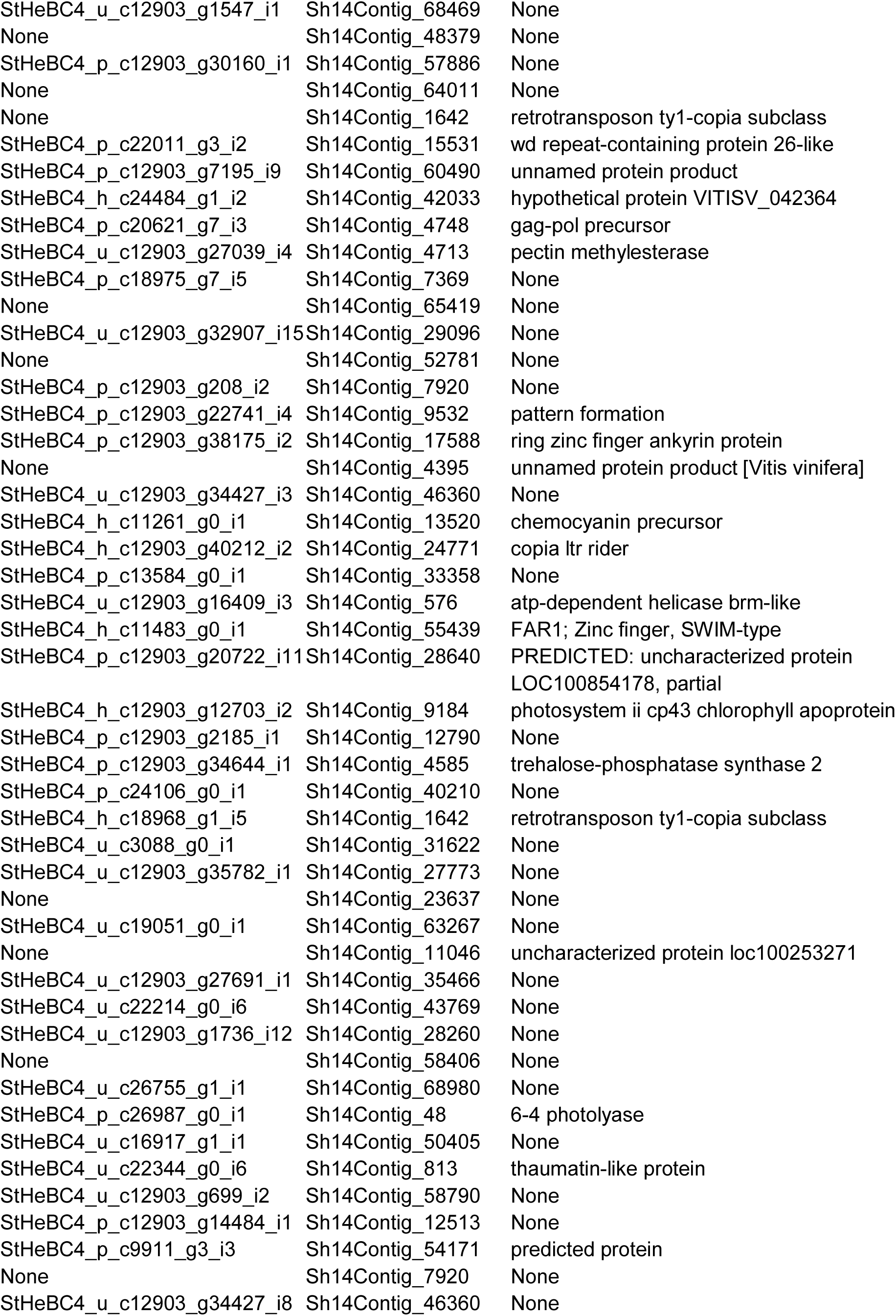
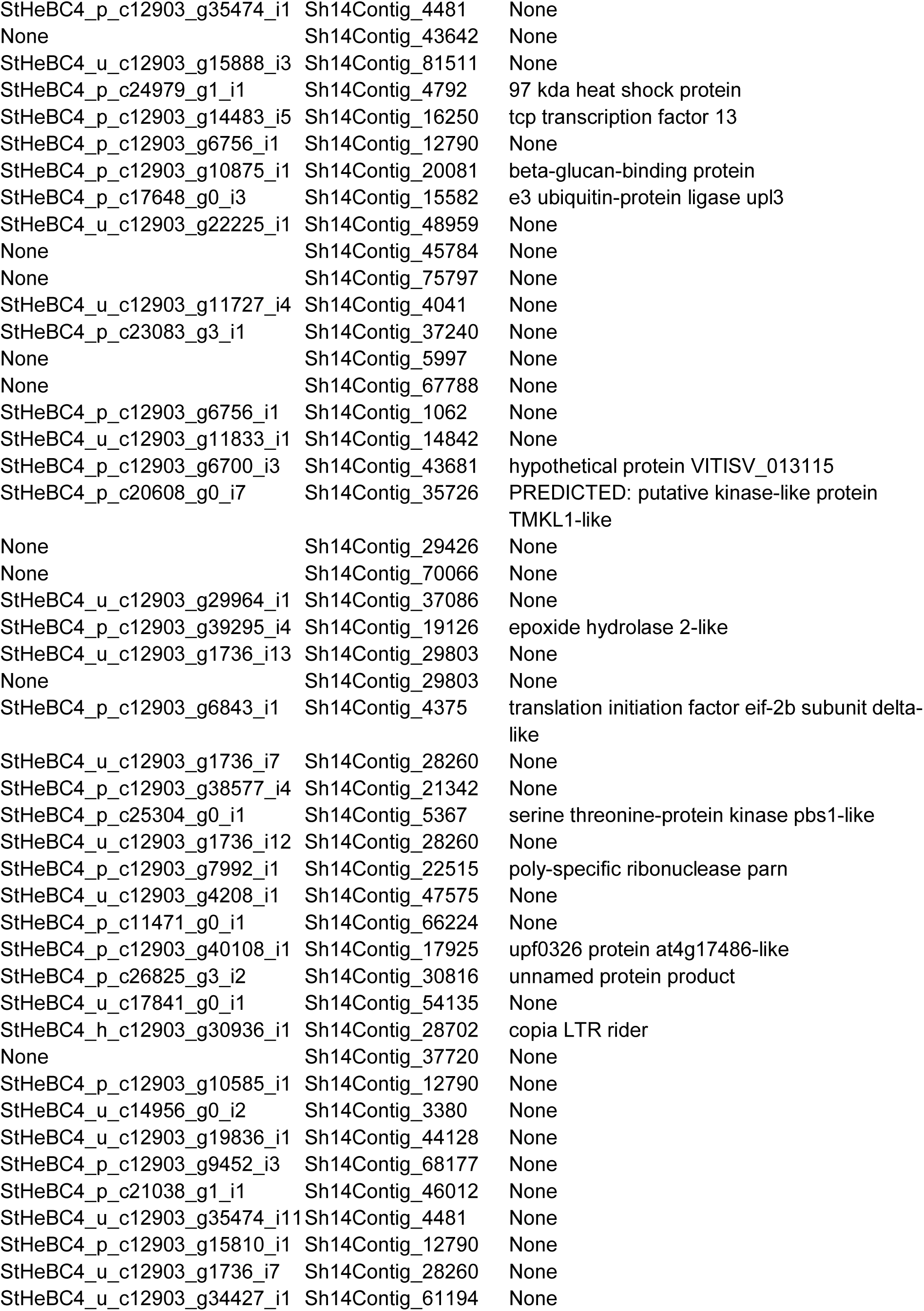
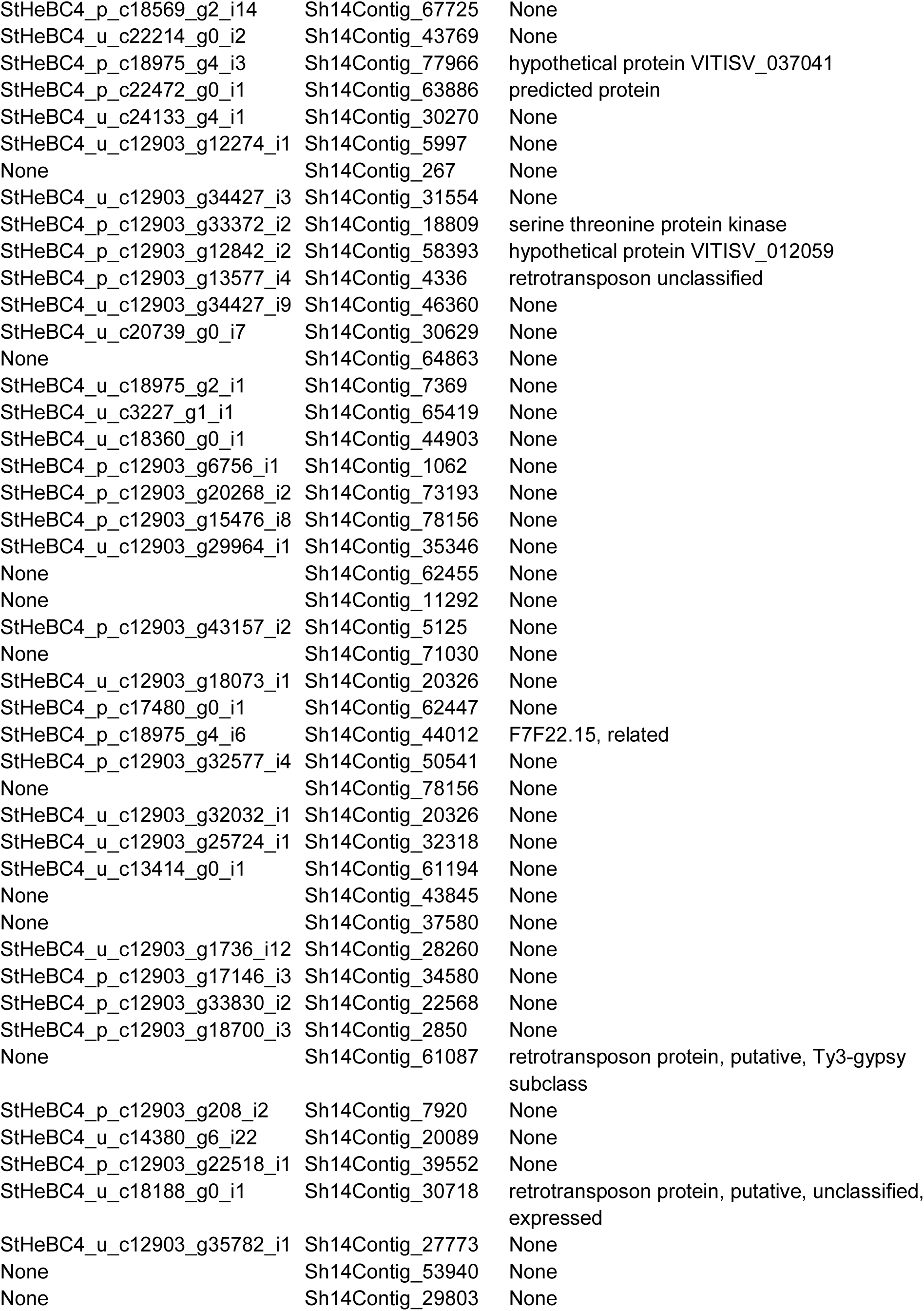
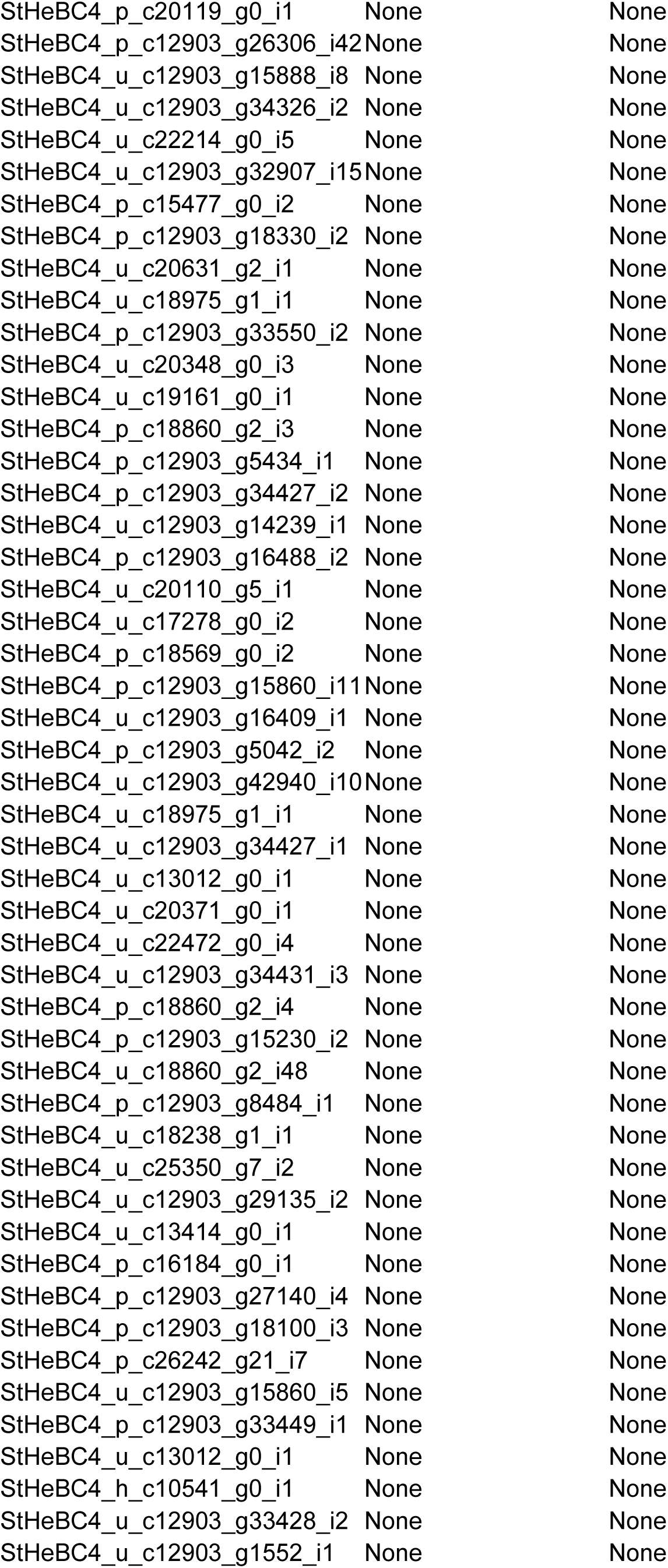
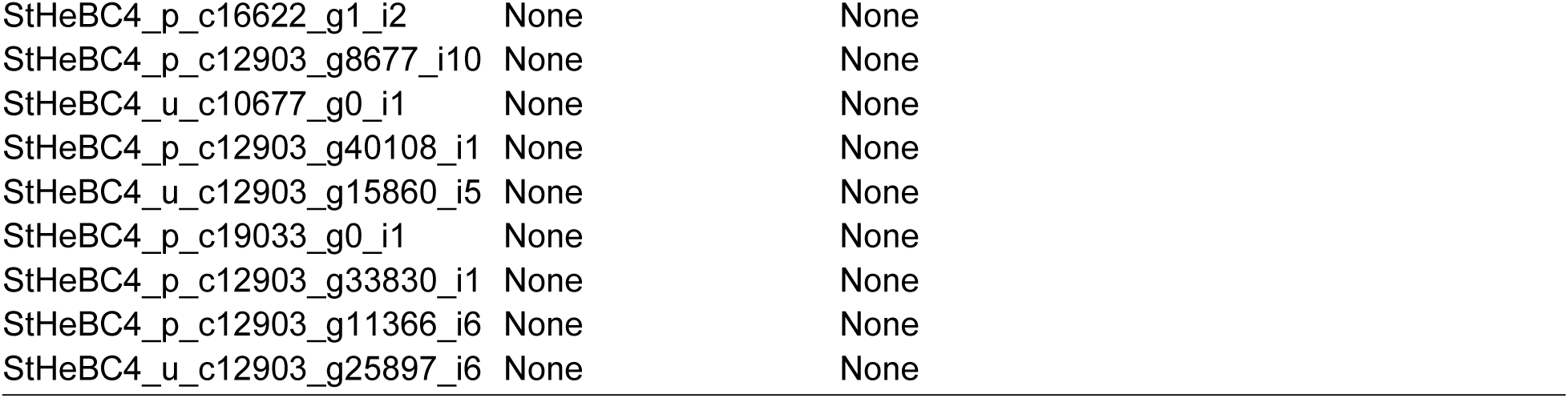
Hits to transcripts from the Parasitic Plant Genome Project (PPGPII) or Sh14v2 (from Yoshida 2019) for contigs assembled from host-associated *k-*mers for *Striga hermonthica* Kisii population (maize vs. finger millet hosts). Transcripts with greater than 85% identity over at least 45 bp are shown.

**Table S4.**
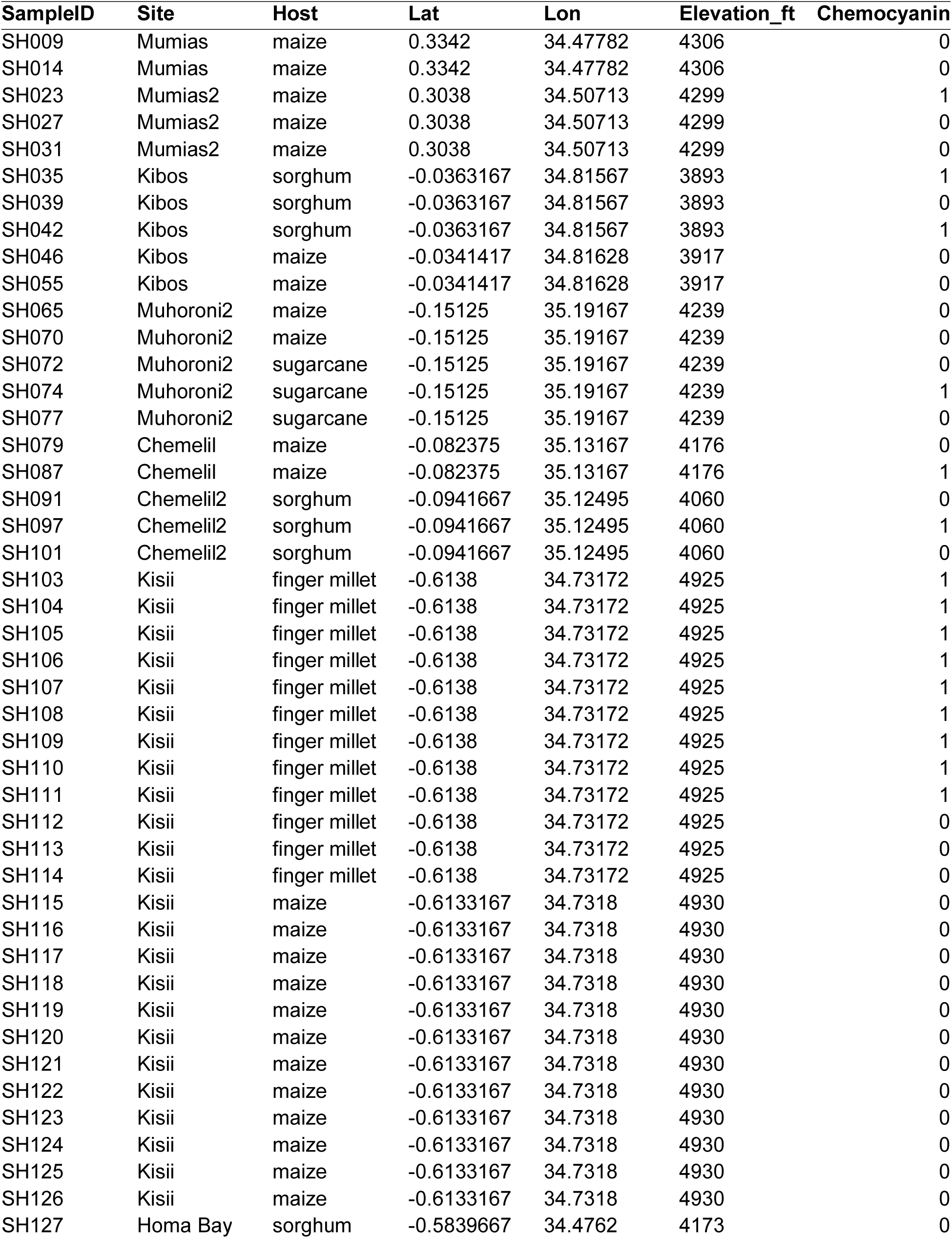
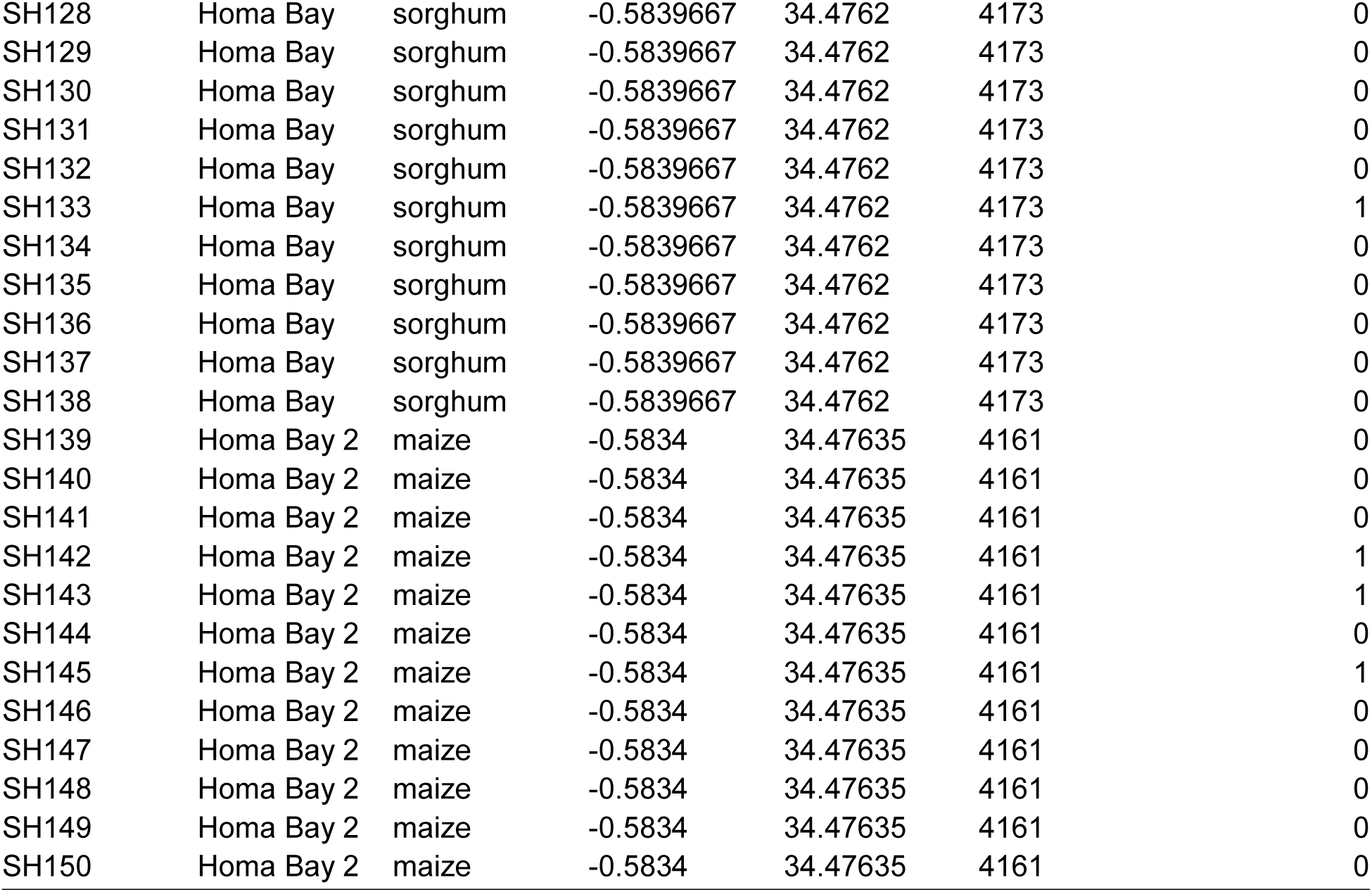
Chemocyanin variant calls for sequenced individuals. Individuals are coded as ‘1’ if they possess the ‘finger millet’ allele or ‘0’ if the allele is missing.

| SampleID | Site | Host | Lat | Lon | Elevation_ft | Chemocyanin |
| --- | --- | --- | --- | --- | --- | --- |
| SH009 | Mumias | maize | 0.3342 | 34.47782 | 4306 | 0 |
| SH014 | Mumias | maize | 0.3342 | 34.47782 | 4306 | 0 |
| SH023 | Mumias2 | maize | 0.3038 | 34.50713 | 4299 | 1 |
| SH027 | Mumias2 | maize | 0.3038 | 34.50713 | 4299 | 0 |
| SH031 | Mumias2 | maize | 0.3038 | 34.50713 | 4299 | 0 |
| SH035 | Kibos | sorghum | -0.0363167 | 34.81567 | 3893 | 1 |
| SH039 | Kibos | sorghum | -0.0363167 | 34.81567 | 3893 | 0 |
| SH042 | Kibos | sorghum | -0.0363167 | 34.81567 | 3893 | 1 |
| SH046 | Kibos | maize | -0.0341417 | 34.81628 | 3917 | 0 |
| SH055 | Kibos | maize | -0.0341417 | 34.81628 | 3917 | 0 |
| SH065 | Muhoroni2 | maize | -0.15125 | 35.19167 | 4239 | 0 |
| SH070 | Muhoroni2 | maize | -0.15125 | 35.19167 | 4239 | 0 |
| SH072 | Muhoroni2 | sugarcane | -0.15125 | 35.19167 | 4239 | 0 |
| SH074 | Muhoroni2 | sugarcane | -0.15125 | 35.19167 | 4239 | 1 |
| SH077 | Muhoroni2 | sugarcane | -0.15125 | 35.19167 | 4239 | 0 |
| SH079 | Chemelil | maize | -0.082375 | 35.13167 | 4176 | 0 |
| SH087 | Chemelil | maize | -0.082375 | 35.13167 | 4176 | 1 |
| SH091 | Chemelil2 | sorghum | -0.0941667 | 35.12495 | 4060 | 0 |
| SH097 | Chemelil2 | sorghum | -0.0941667 | 35.12495 | 4060 | 1 |
| SH101 | Chemelil2 | sorghum | -0.0941667 | 35.12495 | 4060 | 0 |
| SH103 | Kisii | finger millet | -0.6138 | 34.73172 | 4925 | 1 |
| SH104 | Kisii | finger millet | -0.6138 | 34.73172 | 4925 | 1 |
| SH105 | Kisii | finger millet | -0.6138 | 34.73172 | 4925 | 1 |
| SH106 | Kisii | finger millet | -0.6138 | 34.73172 | 4925 | 1 |
| SH107 | Kisii | finger millet | -0.6138 | 34.73172 | 4925 | 1 |
| SH108 | Kisii | finger millet | -0.6138 | 34.73172 | 4925 | 1 |
| SH109 | Kisii | finger millet | -0.6138 | 34.73172 | 4925 | 1 |
| SH110 | Kisii | finger millet | -0.6138 | 34.73172 | 4925 | 1 |
| SH111 | Kisii | finger millet | -0.6138 | 34.73172 | 4925 | 1 |
| SH112 | Kisii | finger millet | -0.6138 | 34.73172 | 4925 | 0 |
| SH113 | Kisii | finger millet | -0.6138 | 34.73172 | 4925 | 0 |
| SH114 | Kisii | finger millet | -0.6138 | 34.73172 | 4925 | 0 |
| SH115 | Kisii | maize | -0.6133167 | 34.7318 | 4930 | 0 |
| SH116 | Kisii | maize | -0.6133167 | 34.7318 | 4930 | 0 |
| SH117 | Kisii | maize | -0.6133167 | 34.7318 | 4930 | 0 |
| SH118 | Kisii | maize | -0.6133167 | 34.7318 | 4930 | 0 |
| SH119 | Kisii | maize | -0.6133167 | 34.7318 | 4930 | 0 |
| SH120 | Kisii | maize | -0.6133167 | 34.7318 | 4930 | 0 |
| SH121 | Kisii | maize | -0.6133167 | 34.7318 | 4930 | 0 |
| SH122 | Kisii | maize | -0.6133167 | 34.7318 | 4930 | 0 |
| SH123 | Kisii | maize | -0.6133167 | 34.7318 | 4930 | 0 |
| SH124 | Kisii | maize | -0.6133167 | 34.7318 | 4930 | 0 |
| SH125 | Kisii | maize | -0.6133167 | 34.7318 | 4930 | 0 |
| SH126 | Kisii | maize | -0.6133167 | 34.7318 | 4930 | 0 |
| SH127 | Homa Bay | sorghum | -0.5839667 | 34.4762 | 4173 | 0 |
| SH128 | Homa Bay | sorghum | -0.5839667 | 34.4762 | 4173 | 0 |
| SH129 | Homa Bay | sorghum | -0.5839667 | 34.4762 | 4173 | 0 |
| SH130 | Homa Bay | sorghum | -0.5839667 | 34.4762 | 4173 | 0 |
| SH131 | Homa Bay | sorghum | -0.5839667 | 34.4762 | 4173 | 0 |
| SH132 | Homa Bay | sorghum | -0.5839667 | 34.4762 | 4173 | 0 |
| SH133 | Homa Bay | sorghum | -0.5839667 | 34.4762 | 4173 | 1 |
| SH134 | Homa Bay | sorghum | -0.5839667 | 34.4762 | 4173 | 0 |
| SH135 | Homa Bay | sorghum | -0.5839667 | 34.4762 | 4173 | 0 |
| SH136 | Homa Bay | sorghum | -0.5839667 | 34.4762 | 4173 | 0 |
| SH137 | Homa Bay | sorghum | -0.5839667 | 34.4762 | 4173 | 0 |
| SH138 | Homa Bay | sorghum | -0.5839667 | 34.4762 | 4173 | 0 |
| SH139 | Homa Bay 2 | maize | -0.5834 | 34.47635 | 4161 | 0 |
| SH140 | Homa Bay 2 | maize | -0.5834 | 34.47635 | 4161 | 0 |
| SH141 | Homa Bay 2 | maize | -0.5834 | 34.47635 | 4161 | 0 |
| SH142 | Homa Bay 2 | maize | -0.5834 | 34.47635 | 4161 | 1 |
| SH143 | Homa Bay 2 | maize | -0.5834 | 34.47635 | 4161 | 1 |
| SH144 | Homa Bay 2 | maize | -0.5834 | 34.47635 | 4161 | 0 |
| SH145 | Homa Bay 2 | maize | -0.5834 | 34.47635 | 4161 | 1 |
| SH146 | Homa Bay 2 | maize | -0.5834 | 34.47635 | 4161 | 0 |
| SH147 | Homa Bay 2 | maize | -0.5834 | 34.47635 | 4161 | 0 |
| SH148 | Homa Bay 2 | maize | -0.5834 | 34.47635 | 4161 | 0 |
| SH149 | Homa Bay 2 | maize | -0.5834 | 34.47635 | 4161 | 0 |
| SH150 | Homa Bay 2 | maize | -0.5834 | 34.47635 | 4161 | 0 |

